# Brain injury accelerates the onset of a reversible age-related microglial phenotype associated with hyperphagocytosis and inflammatory neurodegeneration

**DOI:** 10.1101/2022.05.24.493292

**Authors:** Rodney M. Ritzel, Yun Li, Yun Jiao, Zhuofan Lei, Sarah J. Doran, Junyun He, Rami A. Shahror, Rebecca J. Henry, Shaolin Liu, Bogdan A. Stoica, Alan I. Faden, Gregory Szeto, David J. Loane, Junfang Wu

## Abstract

Lipofuscin is an autofluorescent (AF) pigment formed by lipids and misfolded proteins that accumulates in post-mitotic cells with advanced age. Here we immunophenotyped microglia in the brain of old C57BL/6 mice (>18 months-old) and demonstrate that in comparison to young mice, one third of old microglia are AF, characterized by profound changes in lipid and iron content, phagocytic activity, and oxidative stress. Pharmacological depletion of microglia in old mice eliminated the AF microglia following repopulation and reversed microglial dysfunction. Age-related neurological deficits and neurodegeneration after traumatic brain injury (TBI) were attenuated in old mice lacking AF microglia. Furthermore, hyperphagocytic activity and lipid accumulation in microglia persisted for up to one year after TBI, were modified by *Apoe4* genotype, and chronically driven by phagocyte-mediated oxidative stress. Thus, AF may reflect a pathological state in aging microglia associated with hyperphagocytosis and inflammatory neurodegeneration that can be further accelerated by TBI.

**Teaser:** Traumatic brain injury accelerates age-related pathological phagocytosis and lipofuscin formation in microglia.

## Introduction

Microglia have an exquisite ability to react and respond to environmental changes and stimuli to maintain neuronal homeostasis. As we age, the accumulation of a lifetime of intrinsic and extrinsic stressors gradually increases microglial dysfunction that negatively impacts neurological function. Phenotypically, the changes that occur in post-mitotic microglia with advanced age include increased cytokine and reactive oxygen species (ROS) production, impaired autophagic and metabolic function, and dysregulated phagocytic behavior, which may be further exaggerated in response to injury or disease (*1–3*). One cellular feature, lipofuscin (LF), may reflect dysfunction in all these respective pathways (*4–6*). LF is an autofluorescent (AF) lipopigment formed by lipids, metals, and misfolded proteins (*7*). Although LF granules and associated AF have long been considered a secondary consequence of the brain aging or neurodegenerative disease, there is a mounting evidence that LF may precipitate these changes and play a more active role in neurodegeneration (*8*).

Microglia are known to be relatively long-lived cells that show gradual turnover with limited self-renewal capacity in the aging brain (*9, 10*). This low rate of homeostatic cell division may prevent the dilution of LF aggregates within the aging microglia population. In fact, the buildup of LF in microglia is itself considered a marker of proliferative senescence (*3, 11, 12*); however, causal mechanisms have yet to be established. Microglial surveillance is vital for neuronal health and is closely linked to formation of LF through homeostatic phagocytosis of environmental debris, apoptotic cells, and excess synapses (*7, 13*). However, pathological phagocytosis of neurons and myelin by microglia is also associated with brain damage (*14, 15*); yet it is unclear whether the chronic progression of neurodegeneration and/or white matter disease caused by traumatic brain injury (TBI) are driven by hyperphagocytic activity. Whereas numerous studies have shown that LF and AF in microglia are dramatically increased with normal aging (*16*), the functional relationship between lipid-laden microglia in age-related neurological dysfunction and brain injury has yet to be described. Here we utilize flow cytometry and a simple gating strategy to identify AF microglia in the aging brain that defines hallmark phenotypic features, functional properties, and transcriptomic signatures of this unique and dysfunctional microglial population. Importantly, we demonstrate that pharmacological elimination of AF microglia results in partial reversal of age-related neurological dysfunction and TBI severity. We also demonstrate that moderate-level TBI increases long-term pathological phagocytosis of myelinated neurons, accelerating lipid accumulation and associated inflammatory pathology. These features are modified by *Apoe4* genotype and mediated by phagocyte-mediated oxidative stress mechanisms late after TBI.

## Results

### Age-related autofluorescent microglia have a unique functional phenotype associated with hyperphagocytic activity

We first identified the functional phenotype of naturally occurring AF microglia (see **Supplemental Fig. 3** for gating strategy). Young adult male C57BL/6 mice (3-month-old) were used as a reference to identify AF^lo^ and AF^hi^ microglial populations in aged 18-month-old male C57BL/6 mice by flow cytometry (Fig. 1A). Approximately one-third of microglia in aged mice were AF^hi^ (Fig. 1A, B). Older AF^lo^ microglia had equivalent levels of AF as younger microglia, whereas there was a greater than ∼3 fold increase in AF in old AF^hi^ microglia (Fig. 1C). Compared to young and old bulk populations, old AF^hi^ microglia exhibited signs of hypertrophy and increased granularity as indicated by forward and side scatter properties (**Supplemental Fig. 4A-B**). To confirm whether AF could be attributed to LF, we examined intracellular lipid content, lipid peroxidation, and iron accumulation. Using either BODIPY493/503 or Lipi-Blue probes to measure neutral lipids, we found that old AF^hi^ microglia contain significantly high levels of lipid (Fig. 1D**, Supplemental Fig. 3C**). Lipid peroxidation was increased in old AF^hi^ microglia relative to the AF^lo^ subset as evidenced by a lower reduced:oxidized ratio (Fig. 1E), whereas iron content was dramatically higher in the old AF^hi^ subset (Fig. 1F).

**Figure 1.**
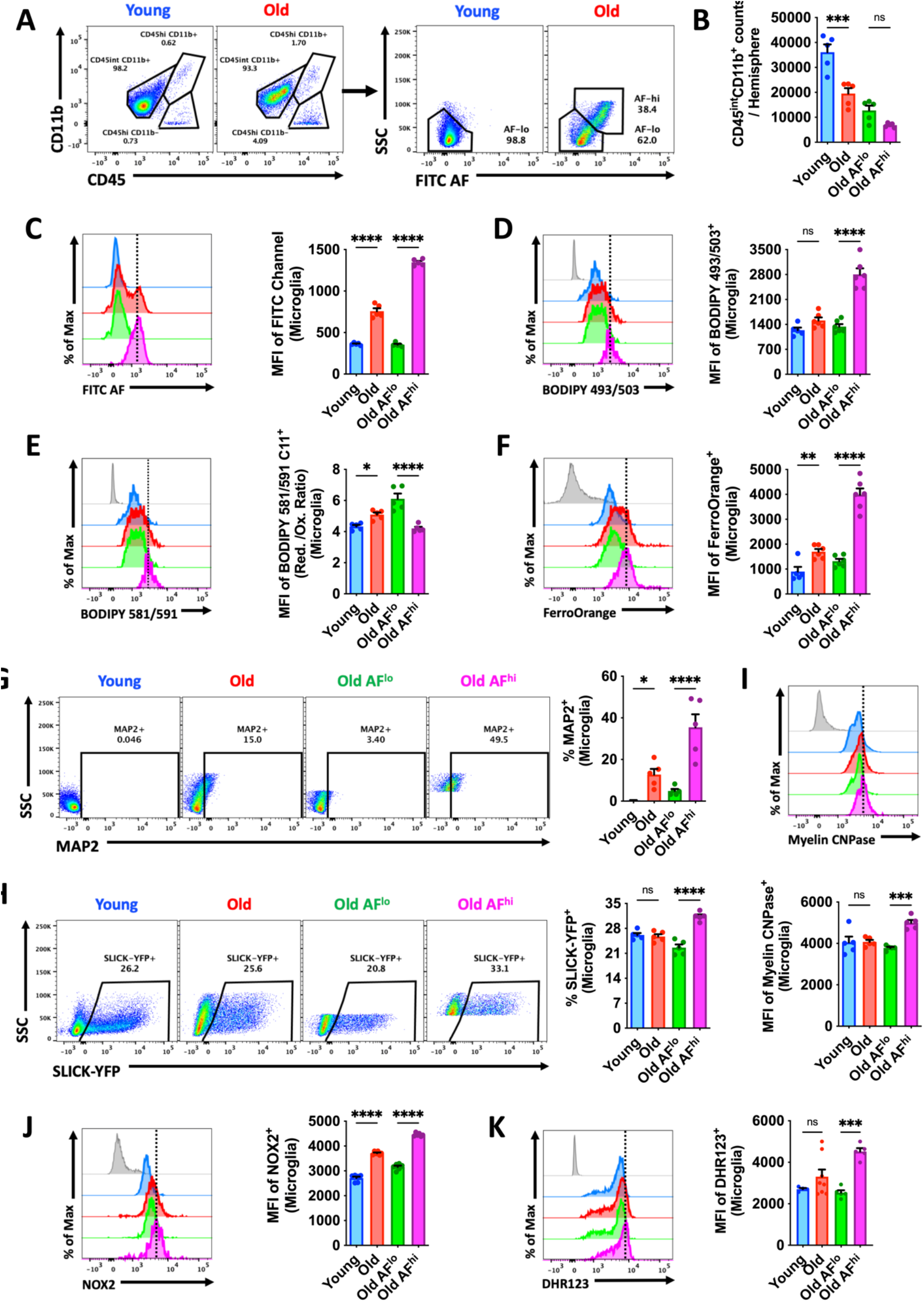
Functional characterization of age-associated AF microglia identifies a lipofuscin phenotype that is hyperphagocytic. **(A)** Representative dot plots showing CD45^int^CD11b^+^ microglia with AF low (AF^lo^) and AF high (AF^hi^) properties in young and old mice, respectively. **(B)** Microglia counts in young and old brain hemispheres, including for each AF subset, is shown. **(C)** The level of AF in each bulk microglial population and subset is quantified using the MFI of the empty FITC channel. Representative histograms are shown next to the relative MFI quantification of **(D)** lipid content, **(E)** lipid peroxidation levels, and **(F)** iron accumulation in microglia as demonstrated using fluorogenic dyes. Oxidative stress was measured by **(G)** intracellular protein expression of NOX2 and **(H)** production of reactive oxygen species using DCF probe. Phagocytic activity was measured by **(I)** intracellular detection of the neuronal marker MAP2, **(J)** fluorescence of neuronal SLICK-YFP reporter mice, and **(K)** intracellular detection of the myelin marker, myelin CNPase. For all experiments, N=5/group. For all flow cytometry histograms, fluorescence minus one (FMO) controls are shown in gray, while microglial populations are color coded according to bar graph. A vertical fiducial line is included for reference. Abbreviations: AF autofluorescence, FITC fluorescein isothiocyanate, Max Maximum, MFI mean fluorescence intensity, Ox oxidation, Red reduction, SSC side scatter, YFP yellow fluorescent protein. Data were analyzed using one-way ANOVA with Bonferroni post-hoc correction for multiple comparisons (*p<0.01, **p<0.01, ***p<0.001, and ****p<0.0001).

To determine whether age-related AF is associated with homeostatic increases in phagocytosis we examined passive and active measures of phagocytic activity. Intracellular detection of the neuronal marker microtubule-associated protein 2 (MAP2) in microglia revealed higher levels in old AF^hi^ relative to AF^lo^ (Fig. 1G). Moreover, compared to the AF^lo^ subset, a significantly higher percentage of AF^hi^ microglia engulfed fluorescently labeled (i.e., Thy1-GFP reporter) apoptotic feeder neurons by *ex vivo* assay (Fig. 1H). Intracellular detection of the myelin antigen, myelin CNPase, confirmed these findings (Fig. 1I). Consistent with the increased ingestion of cargo, LC3II^+^ autophagosome formation was significantly greater in old AF^hi^ microglia (**Supplemental Fig. 3D**). These data are consistent with the hypothesis that age-related microglial AF is associated with LF accumulation, which is either attributable to, or accelerating, the engulfment of neuronal and myelin debris.

LF accumulation is cytotoxic and can potentiate inflammation and oxidative stress (*7*). Old AF^hi^ microglia had higher protein expression of the phagocyte NADPH oxidase (NOX2) (Fig. 1J) and higher ROS production, as measured by DHR123 (Fig. 1K). Intracellular cytokine production of the pro-inflammatory mediator TNF was also higher in AF^hi^ microglia (**Supplemental Fig. 3E**), confirming the association between AF, LF, and inflammatory activity. Old AF^hi^ microglia also had higher glucose demands, increased mitochondrial membrane potential, and lower cytosolic pH (**Supplemental Fig. 3F-H**), which indicates that naturally occurring age-associated AF microglia have an altered metabolic profile.

### Microglia depletion and repopulation reverses the age-related AF phenotype and transcriptomic signature

LF is undegradable but can be efficiently diluted during cell division (*17*). Previous work demonstrated that forced microglial turnover in aged mice can reduce histological markers of LF content (*18*). To expand on this finding, we followed a similar microglial depletion/repopulation protocol by administering the colony-stimulating factor 1 receptor (CSF1R) antagonist, PLX5622, to 18 month-old male C57BL/6 mice in chow for 3 weeks, followed by 4 weeks of repopulation (Fig. 2A-B). No statistically significant differences were seen in the number of brain-resident microglia in the aged brain after repopulation (Fig. 2C). PLX5622-mediated depletion and repopulation eliminated the AF^hi^ subset in old mice, as evidenced by decreased cellular AF, granularity/size, and cytokine and ROS production (Fig. 2D**, Supplemental Fig. 2**).

**Figure 2.**
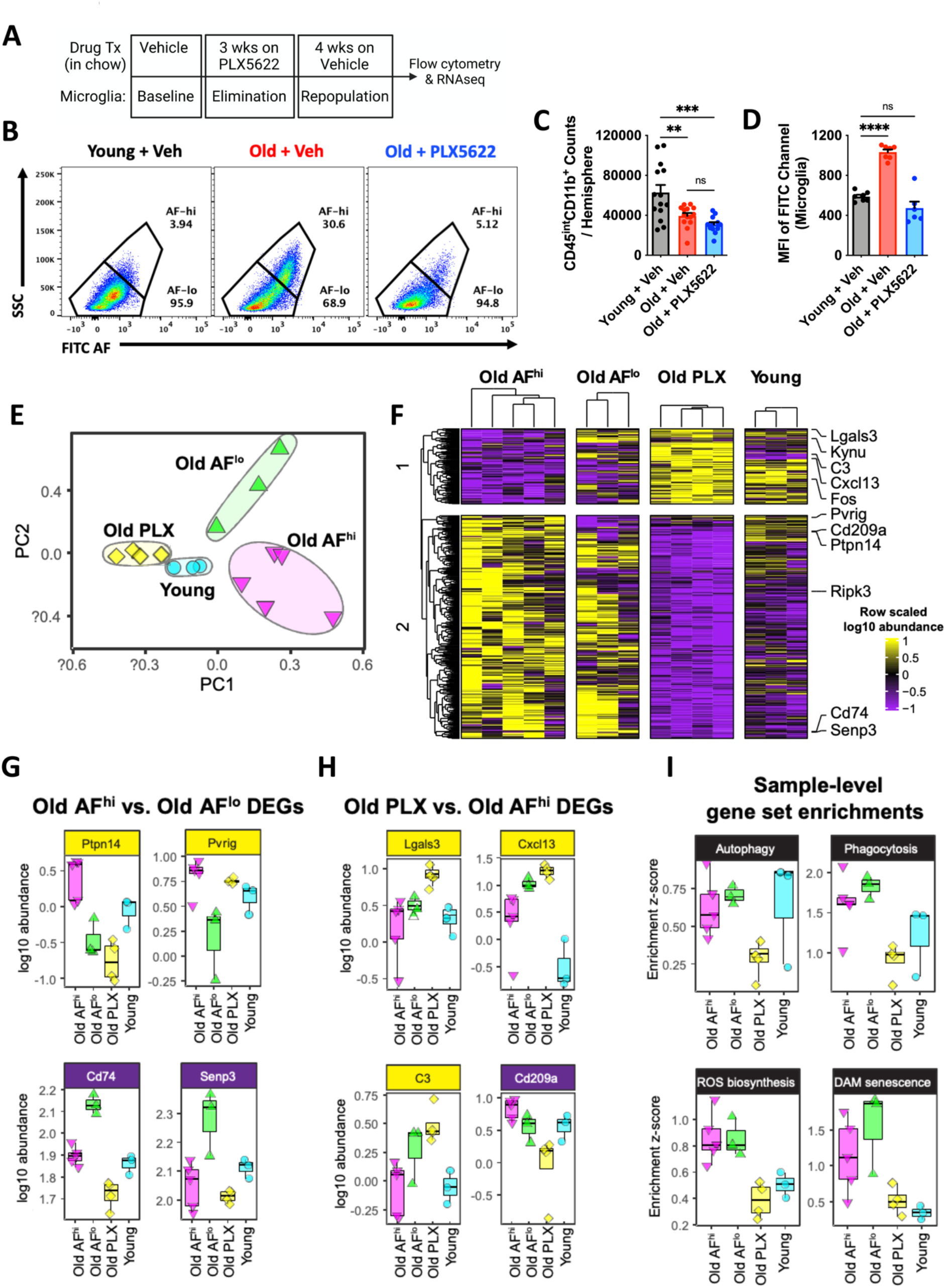
Transcriptomic signature of old, autofluorescent microglia is partially rejuvenated following elimination and four weeks of repopulation. **(A)** Timeline of experimental design for the acute four-week microglial repopulation period is shown. **(B)** A representative dot plot depicts the subsets and group conditions of FACS sorted microglia that were evaluated downstream by RNA sequencing. After four weeks of repopulation, the (**C**) number of microglia and (**D**) mean fluorescence intensity in the FITC channel (i.e., relative autofluorescence) was measured. **(E)** Principal component analysis (PCA) plot of whole transcriptomes for each microglial subset/group is shown. **(F)** Heatmap showing differentially expressed genes (DEGs) comparing old AF^hi^ vs old AF^lo^, or old PLX vs old AF^hi^. Expression is shown as row-scaled log10 abundance. Selected top-ranked DEGs from **(G)** old AF^hi^ vs old AF^lo^, and **(H)** old PLX vs old AF^hi^ comparisons. **(I)** Expression of gene sets for key functional pathways were analyzed using sample-level enrichment analysis (SLEA). Samples were z-scored based on expression of selected gene sets compared to a sample-specific null distribution. Data are represented from two independent experiments (N=6-8 mice/group). Abbreviations: AF autofluorescent, DAM disease-associated microglia, DEGs differentially expressed genes, FITC fluorescein isothiocyanate, hi high, lo low, ns not significant, PC principal component, PLX PLX5622, ROS reactive oxygen species, ssc side scatter, veh vehicle. Data **(C-D)** were analyzed using one-way ANOVA with Bonferroni post-hoc correction for multiple comparisons (**p<0.01, ***p<0.001, and ****p<0.0001).

To characterize the transcriptomic signature of AF subsets, RNA sequencing was performed on FACS-sorted microglia. PCA on whole transcriptomes revealed each microglial subset was distinct along the first two principal components, with old, repopulated microglia clustering closely to young microglia on PC1 and PC2 (Fig. 2E). In contrast, old microglia subsets exhibited significantly more heterogeneity as seen by their intragroup spread along both PC1 and PC2. PC2 also predominantly separated AF^hi^ and AF^lo^ microglia. To define the transcriptional signature of old AF^hi^ microglia, we identified differentially expressed genes between old AF^hi^ versus old AF^lo^, or old PLX5622 versus old AF^hi^ microglia. Hierarchical clustering of differentially expressed genes (DEGs) underscored that old PLX5622 microglia had decreased expression of a gene module associated with old AF^hi^ and old AF^lo^ microglia (Fig. 2F). A module of DEGs was also identified as upregulated in old PLX5622 microglia, downregulated in old AF^hi^, and mixed expression in young. The AF^hi^ gene signature in old mice included up-regulation of *Pvrig*, an immune checkpoint receptor (*19*) and Alzheimer’s disease risk enhancer in macrophages (*20*), and *Ptpn14*, a pro-inflammatory suppressor of SOCS7 (*21*), and down-regulation of *Cd74*, *Senp3, and Kynu*. We tested whether transcriptional signatures could explain functional differences in old AF^hi^ microglia by performing sample-level enrichment analysis (SLEA) on gene sets for phagocytosis, ROS biosynthesis, autophagy, and DAM/senescence pathways (Fig. 2I). The autophagy pathway was decreased in old PLX5622 microglia, but similar in old and young microglia. Phagocytosis, ROS biosynthesis, and DAM senescence pathways were more highly expressed in old AF^hi^ and AF^low^ microglial populations, and attenuated in old PLX5622 microglia. Gene expression for these functional pathways was not significantly different between old AF^hi^ and old AF^lo^ microglia. These findings corroborate earlier work (*22*) and our prior immunophenotyping results (Fig. 1), but also suggest that some functional differences between old AF subsets are not transcriptionally regulated.

### Forced turnover of microglia in old mice restores motor function and cognitive performance

Given that depletion and repopulation of old microglia reversed several features associated with the AF phenotype and reprogrammed transcriptional networks, we next investigated whether forced microglial turnover could modify age-related neurological dysfunction using a diverse battery of behavioral tests (Fig. 3A). At four weeks after repopulation, significant gains in forelimb grip strength were seen in old PLX5622-treated mice (Fig. 3B, Repop group). Age-related deficits in motor coordination using a rotarod test were apparent in old mice, but there was no effect of repopulation on overall performance at either endpoint (Fig. 3C), consistent with earlier findings (*22*). Next, we evaluated gait dynamics and microglial repopulation partially reversed age-related deficits in Regularity Index, Stand, Stride Length, and Body Speed parameters (Fig. 3D-H, Repop group). There were no effects on Swing Speed or Step Cycle (**Supplemental Fig. 5A-B**). Because cerebellar dysfunction can be an underlying cause of uncoordinated forelimb movement, we also examined the age-related cytokine milieu in the cerebellum. Significant reductions in IL-4, IL-9, IL-10, VEGF, M-CSF, IP-10, MIP-1a, MIP-1b, and TNF were found after microglial repopulation in old mice **(Supplemental Fig. 6**). These changes in cerebellar cytokine concentrations represented a return to young adult levels. Surprisingly, gains in forelimb gait function persisted for up to twelve weeks after repopulation (Fig. 3I-P) and mirrored those seen in the hindlimbs (**Supplemental Fig. 7**). Taken together, our data suggest that resetting the inflammatory milieu of brain regions involved in motor function can alleviate age-associated deficits in gait performance.

**Figure 3.**
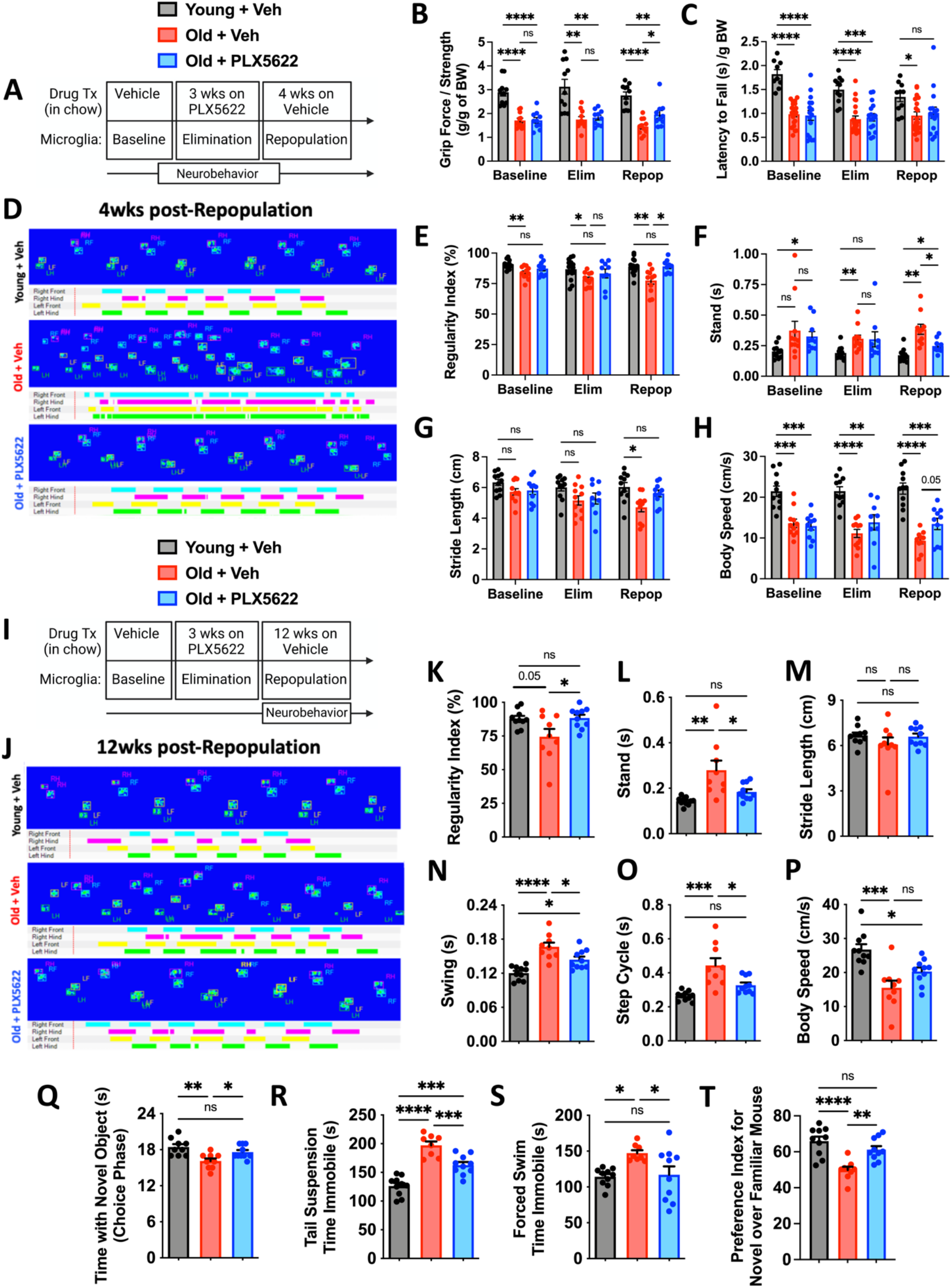
Forced turnover of microglia has lasting beneficial effects on neurological function in aged mice. **(A)** Timeline of experimental design for the acute, four-week microglial repopulation period is shown. **(B)** Forelimb grip strength was quantified for each treatment phase. **(C)** Rotarod performance, as determined by latency to fall, is shown for each phase. **(D)** Representative graphical print view of the CatWalk XT® automated gait analysis system at four weeks after repopulation. Catwalk gait analysis parameters for **(E)** regularity index, **(F)** stand, **(G)** stride length, and **(H)** body speed. **(I)** Timeline of experimental design for the chronic twelve-week microglial repopulation period. **(J)** Representative graphical print view at twelve weeks after repopulation. Catwalk gait analysis parameters for **(K)** regularity index, **(L)** stand, **(M)** stride length, **(N)** swing, **(O)** step cycle, and **(P)** body speed. Data were derived from combined forelimb values for all stand, stride length, swing, and step cycle parameters. **(Q)** Time spent interacting with a novel object during the choice phase of the novel object recognition task at twelve weeks post-repopulation is shown. Time spent immobile during **(R)** tail suspension and **(S)** forced swim tests demonstrate significant reductions in depressive-like behavior at twelve weeks post-repopulation. **(T)** Time spent interacting with a novel mouse during sequence 3 of the social recognition test is shown. For **(B)**, N=10-11/group. For **(C)**, data are combined from two independent experiments, N=11-21/group. For **(E-G)**, N=10-11/group. For **(K-T)**, N=9-10/group. Abbreviations: BW body weight, cm centimeter, Elim elimination, g gram, ns not significant, PLX PLX5622, Repop repopulation, s second, Veh vehicle, wk weeks. Data **(B-C, E-H)** were analyzed using two-way ANOVA with Tukey’s post-hoc correction for multiple comparisons. Data **(K-T)** were analyzed using one-way ANOVA with Bonferroni post-hoc correction for multiple comparisons (*p<0.01, **p<0.01, ***p<0.001, and ****p<0.0001).

To better understand the longer-term benefits of forced turnover in old mice on neurological function, we examined cognitive, socialization, and depression-like behaviors at twelve weeks post-repopulation (Fig. 3I). In cognitive testing, microglial repopulation in old mice increased the interaction with the novel object during the choice phase of the novel object recognition test, indicating that forced turnover of old microglia improved declarative memory (Fig. 3Q). Age-related depression-like behaviors were also reduced following microglial repopulation in old mice as evidenced by increased mobility in both the tail suspension (Fig. 3R) and forced swim tests (Fig. 3S). Furthermore, age-related deficits in social recognition, or preference for social novelty, were partially reversed in old mice with forced microglial turnover (Fig. 3T).

### The beneficial effects of forced turnover of AF microglia are sustained for months and modified by *Apoe4* genotype

We then investigated whether elimination of the AF^hi^ subset was sustained late after repopulation or if the phenotype eventually re-emerges with advanced age (Fig. 4A). Interestingly, age-related reductions in microglia number/yield were partially mitigated in old mice at twelve weeks post-repopulation (Fig. 4B). Young and old repopulated microglia showed stable decreases in cell size and granularity compared to vehicle-treated control groups (**Supplemental Fig. 8A and 8B,** respectively). Importantly, flow cytometry measurements of AF, lipid accumulation, and iron content were each significantly attenuated at 12 weeks post-repopulation, an effect only observed in aged mice (Fig. 4C-F). A significant decrease after repopulation was also seen in microglial production of pro-inflammatory cytokines and ROS, as measured by DCF (Fig. 4G-H and 4I, respectively). Consistent with this change in microglial phenotype, cellular alterations in metabolic function were also stably reversed late after forced turnover (**Supplemental Fig. 8C-E**). These results demonstrate that the beneficial effects of forced repopulation on microglial phenotype are long-lasting.

**Figure 4.**
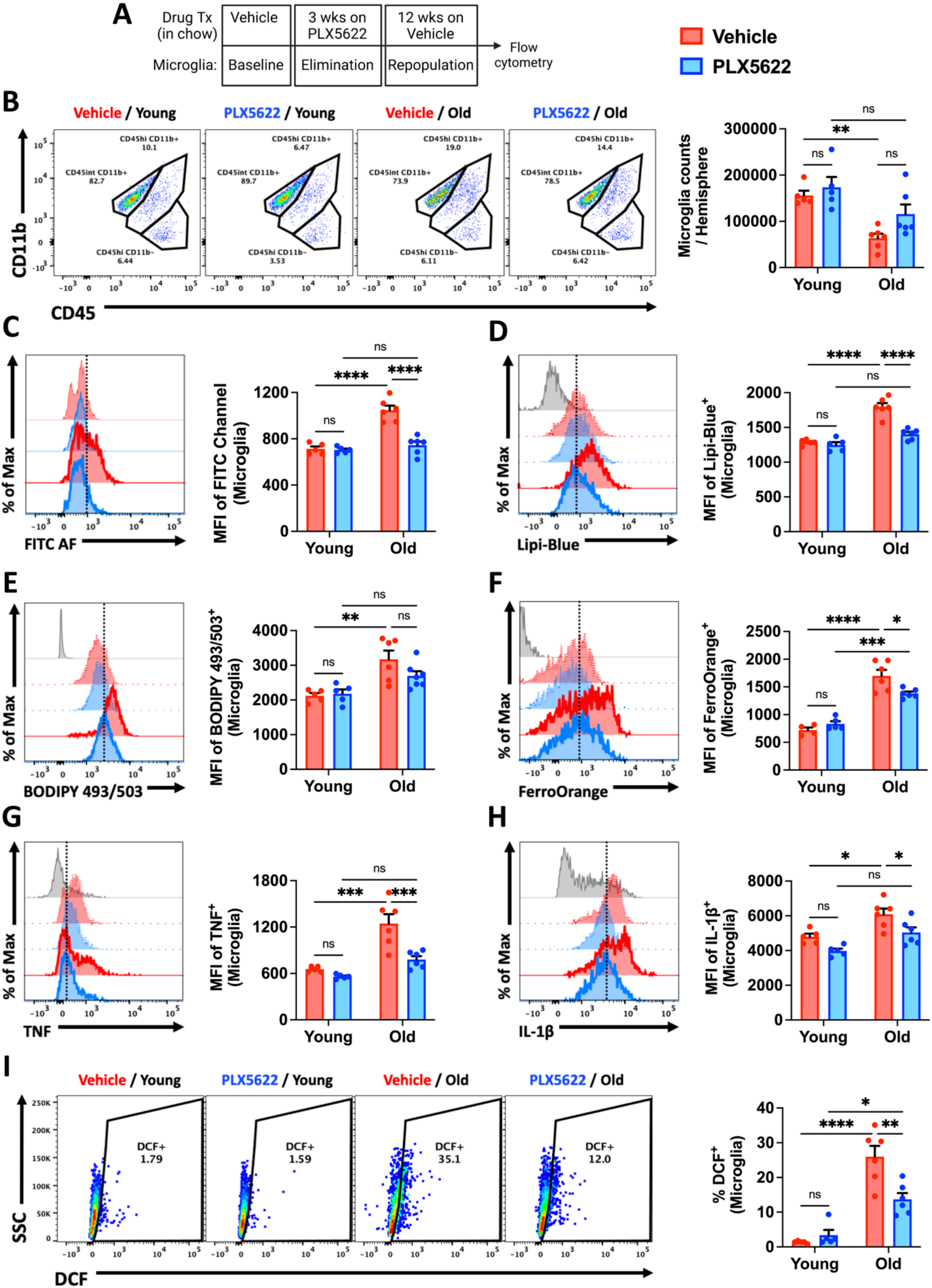
Reductions in AF-related phenotype persist for months after microglial repopulation in aged mice. **(A)** Timeline of experimental design for the chronic, twelve-week microglial repopulation period is shown. **(B)** Representative dot plots of young and old microglia at twelve weeks post-repopulation. Quantification of microglia counts per hemisphere is shown. Representative histograms and quantification of the AF-related phenotype as measured by MFI of **(C)** FITC AF, **(D-E)** lipid content, **(F)** iron accumulation, **(G-H)** pro-inflammatory cytokine production, and **(I)** reactive oxygen species production. PLX5622 treatment (i.e., elimination and repopulation) showed enduring decreases in all AF-related microglial biomarkers. For all experiments, N=4-6/group. For all flow cytometry histograms, fluorescence minus one (FMO) controls are shown in gray, young groups are shown with no outline, old groups are shown with bold outlines, and treatment groups are color coded according to bar graph and figure legend (Vehicle in red, PLX5622 in blue). A vertical fiducial line is included for reference. Abbreviations: AF autofluorescence, FITC fluorescein isothiocyanate, ns not significant, Max Maximum, MFI mean fluorescence intensity, SSC side scatter. Data were analyzed using two-way ANOVA with Tukey post-hoc correction for multiple comparisons (*p<0.01, **p<0.01, ***p<0.001, and ****p<0.0001).

The age-dependent effects of microglial repopulation on the AF phenotype suggest that mechanisms associated with normal aging contribute to its emergence. To test this, we investigated whether old mice harboring the human *apolipoprotein E4* (h*Apoe4*) genotype, the strongest genetic risk for developing late-onset Alzheimer’s disease (AD) (*23*), modified the AF microglial phenotype. Relative to age-matched wildtype controls, microglia from old h*Apoe4*^+/+^ mice (16-20 months-old) exhibited significantly altered numbers, AF, lipid accumulation, and iron content (**Supplemental Fig. 9A-D**). Moreover, protein expression levels of CD68, Lamp1, and Clec7a, which are upregulated in AD and associated with lysosomal-phagocytic and lipid metabolism pathways (*24*), were also elevated in aged h*Apoe4*^+/+^ microglia (**Supplemental Fig. 9E-G**). Interestingly, these changes were associated with a robust increase in expression of the senescence marker p16^ink4a^ (**Supplemental Fig. 9H**). Taken together, these data indicate that *Apoe* genotype can modify severity of the AF phenotype in microglia of older mice.

### Old AF microglia have an exacerbated neuroinflammatory response to TBI

The improvements seen in functional measures that typically decline or become exaggerated with advanced age led us to hypothesize that elimination of AF^hi^ microglia may have a protective effect on the aged brain’s response to injury. To test this hypothesis, we subjected old mice (18 months-old) to moderate-level controlled cortical impact or sham surgery at 4 weeks following microglial repopulation (Fig. 5A). At 3 days post-injury, TBI-induced microglial proliferation and leukocyte extravasation were significantly attenuated in old repopulated mice relative to sham and vehicle controls, as evidenced by lower cell counts (Fig. 5B). TBI-induced production of ROS and IL-1β, including p16^ink4a^ protein expression, were all significantly reduced in old repopulated microglia compared to sham or injured vehicle control (Fig. 5C-E). To measure the impact of TBI on the inflammatory milieu in the perilesional cortex we examined cytokine concentrations using multiplex ELISA. TBI caused an increase in Eotaxin, MIG, MIP-2, IL-1α, G-CSF, and M-CSF concentrations in vehicle, but not PLX5622-treated TBI mice (Fig. 5F). These results suggest that the presence of AF^hi^ microglia exacerbate acute neuroinflammation following TBI in old mice.

**Figure 5.**
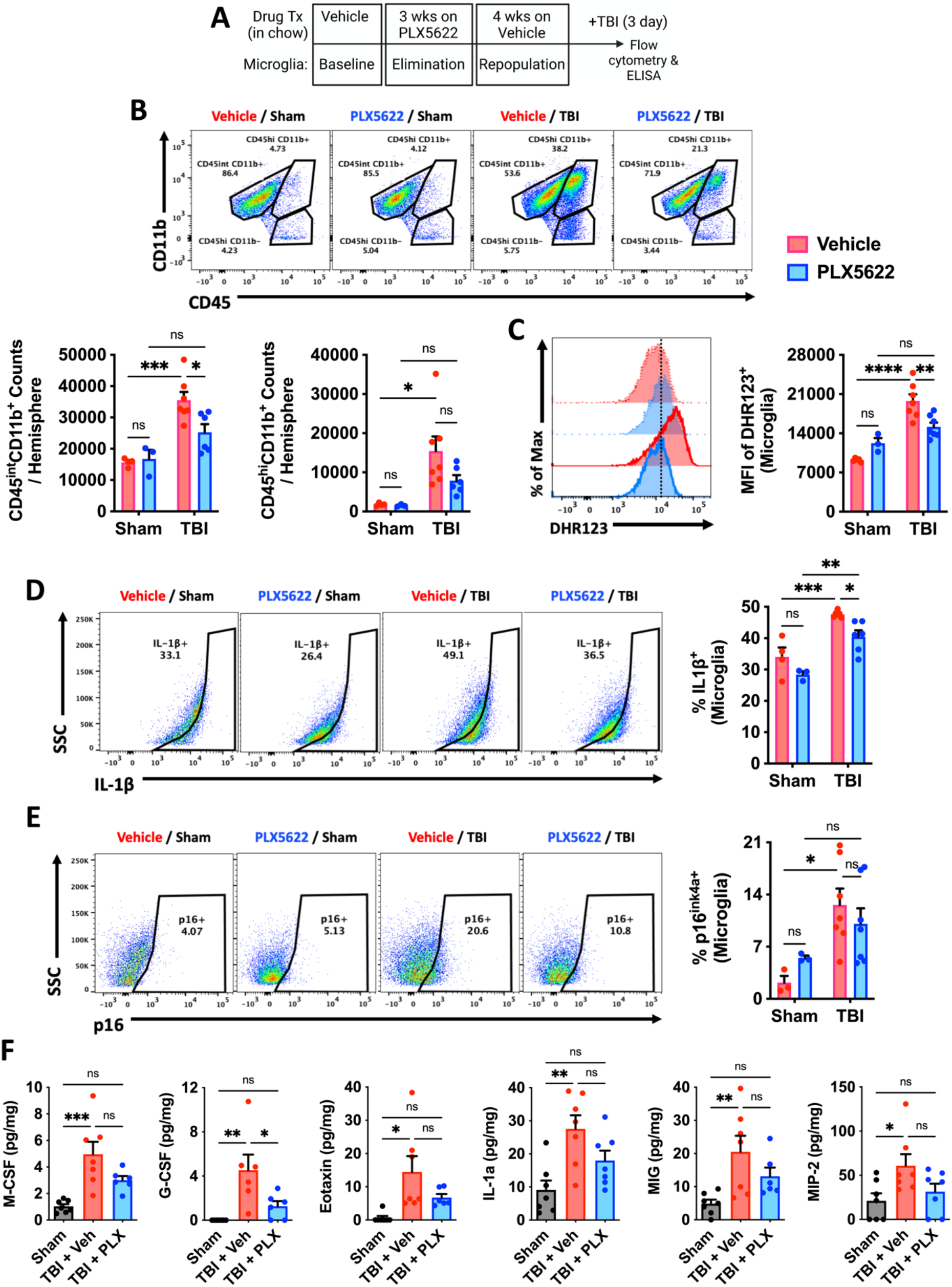
Forced turnover of microglia in aged mice decreases sensitivity to TBI. **(A)** Timeline of experimental design immediately prior to sham and TBI surgery. **(B)** Representative dot plots of immune populations in the ipsilateral brain hemisphere at 3 days after TBI. Quantification of CD45^int^CD11b^+^ microglia and CD45^hi^CD11b^+^ infiltrating myeloid cell counts per hemisphere are shown for aged, surgery and treatment groups. **(C)** A representative histogram of DHR123^+^ microglia is shown next to the relative MFI quantification of reactive oxygen species production. In the associated histogram, young groups are shown with no outline, old groups are shown with bold outlines, and treatment groups are color coded according to bar graph and figure legend (Vehicle in red, PLX5622 in blue). A vertical fiducial line is included for reference. Representative dot plots depicting **(D)** IL1β production and **(E)** p16 expression in microglia are shown next to quantification of cell frequencies. **(F)** Cytokine protein concentrations in the peri-lesional cortex as measured by ELISA. No differences were seen between sham control groups after treatment, and so data for both sham groups were combined. For all cytokines (M-CSF, G-CSF, Eotaxin, IL1a, MIG, and MIP-2), TBI acutely increased concentrations in vehicle but not PLX5622-treated groups. For all experiments, N=3-4/sham and N=6-7/TBI group. Abbreviations: ns not significant, Max Maximum, MFI mean fluorescence intensity, mg milligram, pg picrogram, PLX PLX5622, SSC side scatter, TBI traumatic brain injury, Veh vehicle. Data **(B-E)** were analyzed using two-way ANOVA with Tukey post-hoc correction for multiple comparisons. Data **(D)** were analyzed using one-way ANOVA with Bonferroni post-hoc correction for multiple comparisons (*p<0.01, **p<0.01, ***p<0.001, and ****p<0.0001).

### Elimination of AF microglia in old mice reduces chronic hyperphagocytosis and inflammatory activity following TBI

To see whether attenuated microglial responses to TBI were stable over time, we injured old mice (18 months-old) at 12 weeks following microglial repopulation and evaluated inflammatory cell functions at two weeks post-injury (Fig. 6A). While the number of microglia did not differ significantly between groups after injury, the total number of CD45^hi^ myeloid cells and lymphocytes recruited to the injured, ipsilateral hemisphere was markedly reduced in repopulated old mice (Fig. 6B). TBI-induced expression of the phagocytic marker, CD68, was attenuated in repopulated ipsilateral microglia compared to both contralateral (intact uninjured hemisphere) and injured vehicle controls (Fig. 6C). Moreover, the lipid accumulation and ROS levels seen in the control group after TBI were significantly reduced in repopulated old microglia (Fig. 6D-F). Additional staining confirmed the sustained attenuation of TBI-induced lysosomal-phagocytic functions in repopulated microglia (**Supplemental Fig. 10**). TBI caused an elevation in IP-10 and MIG tissue concentrations in the perilesional cortex that were also reduced in repopulated mice (Fig. 6G). These data suggested that the absence of AF^hi^ microglia in old mice was associated with lower levels of chronic neuroinflammation after TBI.

**Figure 6.**
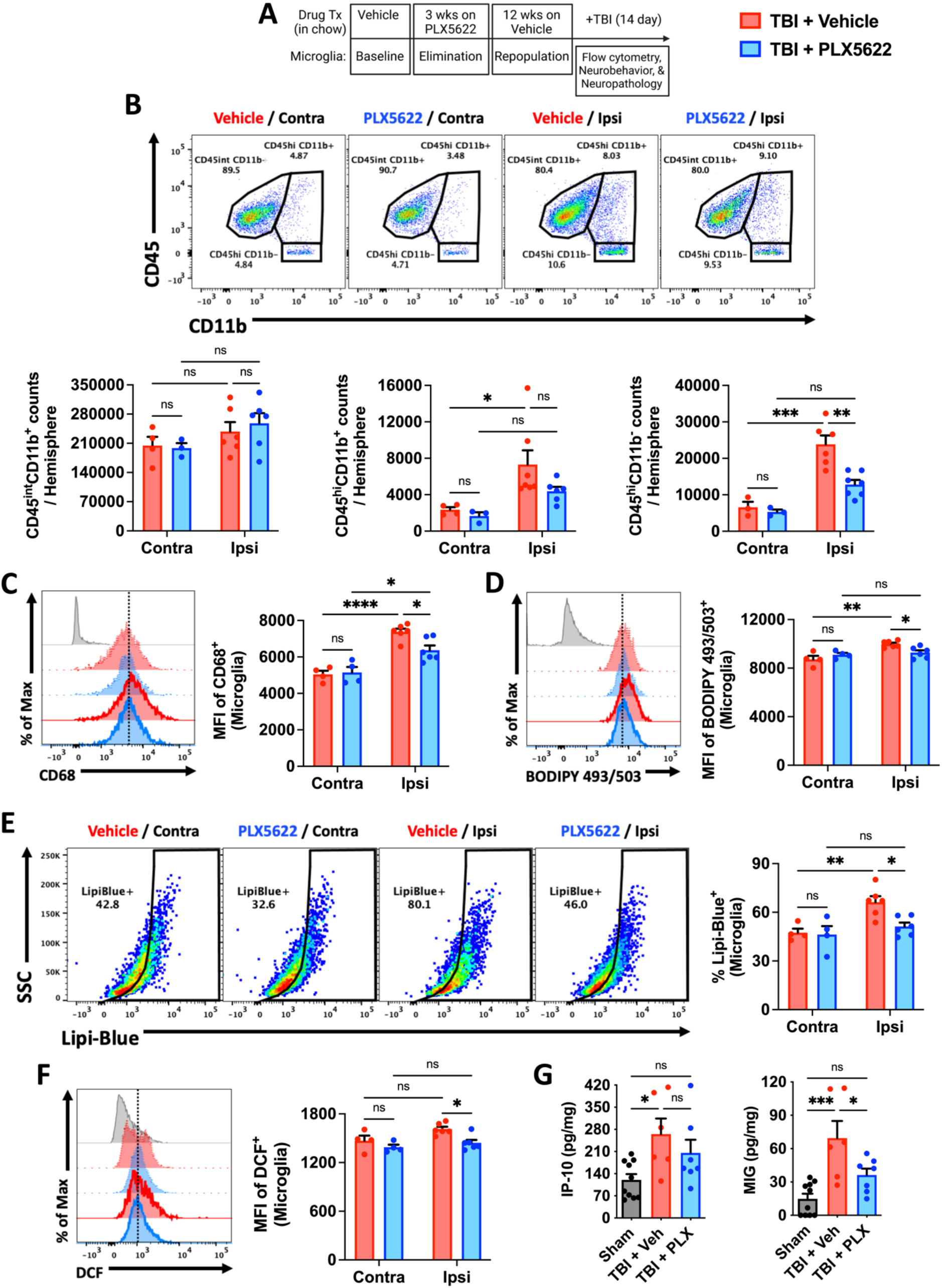
Forced turnover of microglia in aged mice reduces chronic activation and emergence of the AF-related phenotype after TBI. **(A)** Timeline of experimental design immediately prior to sham and TBI surgery. **(B)** Representative dot plots of immune populations in the ipsilateral and contralateral (internal control) brain hemispheres at 2 weeks after TBI. Quantification of CD45^int^CD11b^+^ microglia (left) and infiltrating CD45^hi^CD11b^+^ myeloid (center) and CD45^hi^CD11b^-^ lymphocyte (right) counts per hemisphere are shown for aged, injury and treatment groups. **(C)** A representative histogram of CD68^+^ microglia is shown next to the MFI quantification of this phagocytosis marker. **(D)** A representative histogram shows the relative level of neutral lipids in microglia from each hemisphere after TBI. **(E)** Representative dot plots depict the percentage of Lipi-Blue-positive microglia after TBI. The frequency of lipid droplet-containing microglia is quantified. **(F)** A representative histogram shows the relative level of reactive oxygen species production in microglia as measured by DCF probe. **(G)** Cytokine protein concentrations in the peri-lesional cortex as measured by ELISA. No differences were seen between sham control groups after treatment, and so data for both sham groups were combined. For all cytokines (IP-10 and MIG), TBI chronically increased concentrations in vehicle but not PLX5622-treated TBI groups. For all experiments, N=3-4/contralateral and N=6-7/ipsilateral (i.e., TBI) group. For all flow cytometry histograms, fluorescence minus one (FMO) controls are shown in gray, contralateral groups are shown with no outline, ipsilateral groups are shown with bold outlines, and treatment groups are color coded according to bar graph and figure legend (TBI + Vehicle in red, TBI + PLX5622 in blue). A vertical fiducial line is included for reference. Abbreviations: Contra contralateral, Ipsi ipsilateral, ns not significant, Max Maximum, MFI mean fluorescence intensity, mg milligram, pg picrogram, PLX PLX5622, SSC side scatter, TBI traumatic brain injury, Veh vehicle. Data **(B-F)** were analyzed using two-way ANOVA with Tukey post-hoc correction for multiple comparisons. Data **(G)** were analyzed using one-way ANOVA with Bonferroni post-hoc correction for multiple comparisons (*p<0.01, **p<0.01, ***p<0.001, and ****p<0.0001).

### Elimination of AF microglia in old mice improves neurobehavioral outcomes and reduces chronic neurodegeneration after TBI

To validate our previous findings, we also performed neurobehavioral testing to assess functional outcomes of old mice up to two weeks after TBI (Fig. 6A). Compared to injured vehicle control old mice, repopulated old mice also exhibited significantly better preservation of spatial-working (Fig. 7A) and recognition (Fig. 7B-C) memory in the Y-maze and novel object recognition tests, respectively, indicating that forced turnover of AF microglia can preserve long-term cognitive function after TBI.

**Figure 7.**
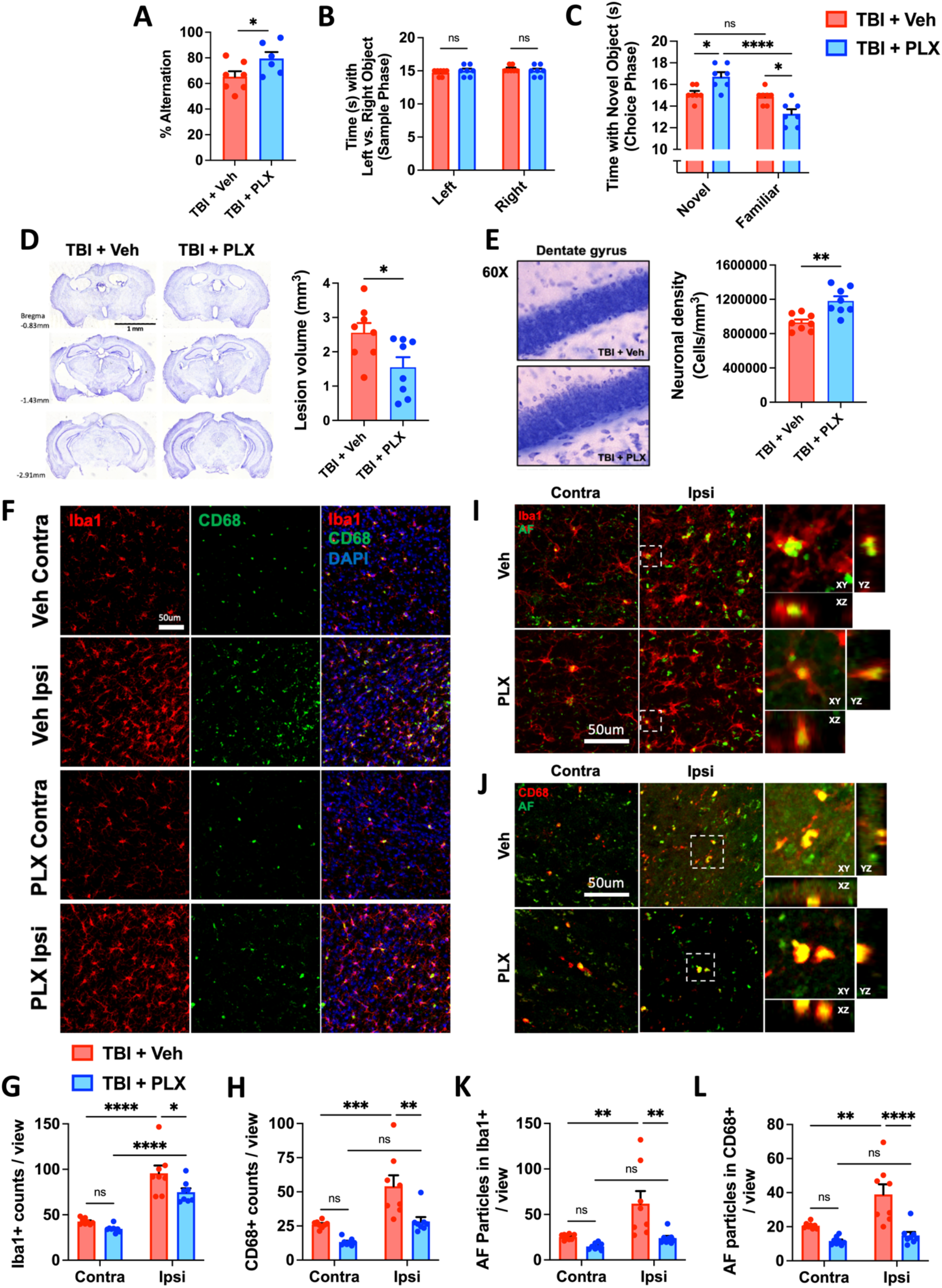
Increased neurological recovery and neuroprotection after TBI following microglial repopulation is associated with attenuation of injury-induced AF phenotype. A behavioral and neuropathological assessment was performed on old TBI mice after a twelve-week repopulation period. **(A)** The percentage of spontaneous alternations in the Y-maze at 7 days post-TBI is shown for each group. **(B)** Time spent interacting with the left and right objects during the sample phase of the novel object recognition test at 7 days post-TBI is shown. **(C)** Time spent interacting with a novel object during the choice phase of the novel object task is shown. **(D)** Histological analysis of brain at 2 weeks post-TBI. Representative images and lesion volume quantification of cresyl violet-stained brain sections demonstrate significantly smaller volumes in the PLX5622-treated group. **(E)** Stereological counts of cresyl violet-stained neurons in the dentate gyrus of the ipsilateral hippocampus. **(F, I, J)** *In situ* analysis of Iba1-positive microglia in the perilesional cortex at two weeks post-TBI. TBI increased the number of **(G)** Iba1-positive microglia/macrophages and **(H)** CD68-positive phagocytes per view. The number of AF particles in **(K)** Iba1-positive and **(L)** CD68-positive cells was quantified. TBI increased the AF particle count in phagocytes of vehicle-but not PLX5622-treated groups. Scale bars, **(F)** 1 mm and **(I-J)** 50 μm. For all behavioral experiments, **(A-C)** N=6-7/group. For all histology experiments, N=8/group. Abbreviations: AF autofluorescent, Contra contralateral, Ipsi ipsilateral, ns not significant, mm milliliter, s second, PLX PLX5622, TBI traumatic brain injury, Veh vehicle. Data **(A, D-E)** were analyzed using Student’s t-test. Data **(B-C, G-L)** were analyzed using two-way ANOVA with Tukey post-hoc correction for multiple comparisons (*p<0.01, **p<0.01, ***p<0.001, and ****p<0.0001).

Next, we performed an unbiased histopathological assessment of microglial activation and neurodegeneration. TBI-induced lesion volumes in old mice were significantly reduced in repopulated mice, as measured using cresyl violet (Fig. 7D). Stereological quantification of surviving neurons in the dentate gyrus region of the hippocampus revealed significantly greater preservation of neurons in the repopulated brain compared to injured vehicle controls (Fig. 7E). These changes were associated with a significant reduction in Iba1-positive microglia/macrophages (Fig. 7F-G) in the injured cortex of old mice. Consistent with our previous findings, we observed reduced numbers of CD68-positive microglia/macrophages and AF phagocytes (Fig. 7H and 7I-L, respectively) in the repopulated brain compared to injured vehicle controls. Together, these data support the notion that old AF^hi^ microglia contribute to age-related neuropathology.

### Hyperphagocytosis and lipid accumulation are hallmarks of post-traumatic chronic microglial activation in young mice, which is mediated by oxidative stress

Our results in old mice implied that TBI can exacerbate the AF phenotype in microglia, characterized by hyperphagocytosis and lipid accumulation. To examine this in more detail, we evaluated microglia along these phenotypic measures at chronic stages of TBI (i.e., one-year post-injury) in young C57BL/6 mice (3-month-old). Microglia from chronically injured mice were significantly more likely to engulf apoptotic feeder neurons (Fig. 8A) and had significantly increased lipid content (Fig. 8B) than age-matched sham or young controls. Further analysis confirmed that microglia engulfment of neurons resulted in increased myelin uptake (Fig. 8C) and expression of CD68 (Fig. 8D). Moreover, microglia which contained high levels of lipid also contained high levels of iron (Fig. 8E), consistent with LF. Thus, the AF microglial immunopathological features are chronically altered after TBI.

**Figure 8.**
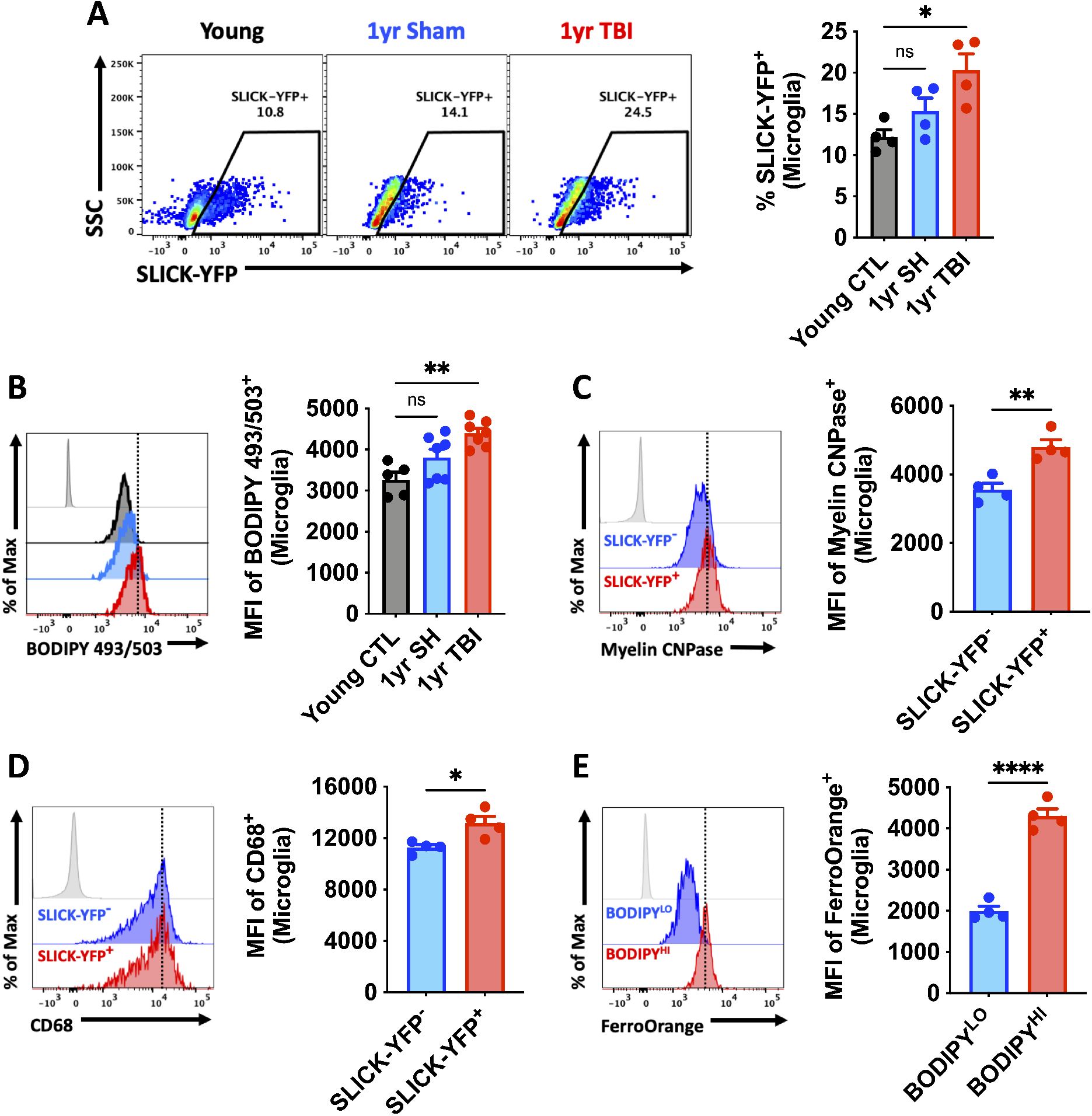
TBI accelerates development of the AF-related phenotype and chronically increases microglial phagocytosis of apoptotic neurons and myelin. Microglial phagocytosis of neurons was assessed by *ex vivo* engulfment of apoptotic cortical neurons isolated from SLICK-YFP transgenic reporter mice. **(A)** Representative dot plots of microglia expressing YFP at one year after TBI. Quantification of phagocytic microglia is shown (right). **(B)** A representative histogram of BODIPY^+^ microglia at one year after TBI is shown next to the MFI quantification of this neutral lipid marker. **(C)** A representative histogram shows the relative protein expression of myelin CNPase in YFP-phagocytic and non-phagocytic microglia at one year post-TBI. **(D)** A representative histogram shows the relative expression level of the phagocytosis marker CD68 in YFP-positive and YFP-negative microglia. The MFI of CD68-positive microglia is quantified. **(E)** A representative histogram shows the relative level of intracellular iron deposition in subsets of microglia that low and high in neutral lipid content at one year after TBI. For all experiments, N=4-7/group. For all histograms **(B-E)**, fluorescence minus one (FMO) controls are shown in light gray, while all other groups are shown with bold outline as color coded according to the bar graph axis labels. A vertical fiducial line is included for reference. Abbreviations: CTL control, Hi high, ns not significant, Lo low, Max Maximum, MFI mean fluorescence intensity, SH sham, SSC side scatter, TBI traumatic brain injury, YFP yellow fluorescent protein. Data **(A-B)** were analyzed using one-way ANOVA with Bonferroni post-hoc correction for multiple comparisons. Data **(C-E)** were analyzed using Student’s t-test (*p<0.01, **p<0.01, and ****p<0.0001).

Finally, we attempted to elucidate an injury-dependent mechanism through which the development of this pathological AF microglial phenotype is accelerated after TBI. Because LF is exacerbated by oxidative stress and given the close association between phagocytosis and ROS production, we utilized transgenic mice which have a genetic deletion of the microglial Hv1 proton channel, which is required for NOX2 activity in phagocytes (*25*). At one-year after TBI, microglia from Hv1-knockout mice failed to show a chronic, injury-related increase in phagocytosis, lipid content, and oxidative stress (Fig. 9A-D). Importantly, Hv1-knockout mice had significantly better functional outcomes, including grip strength (Fig. 9E), social recognition (Fig. 9F-G), and depressive-like behavior (Fig. 9H). In the novel object recogition test, wildtype mice spent considerably less time interacting with the novel object after chronic TBI compared to Hv1-knockout mice (Fig. 9I-J). Together, these data provide proof-of-principle evidence that phagocytic ROS intrinsically drives the emergence of the age-associated and chronic AF phenotype.

**Figure 9.**
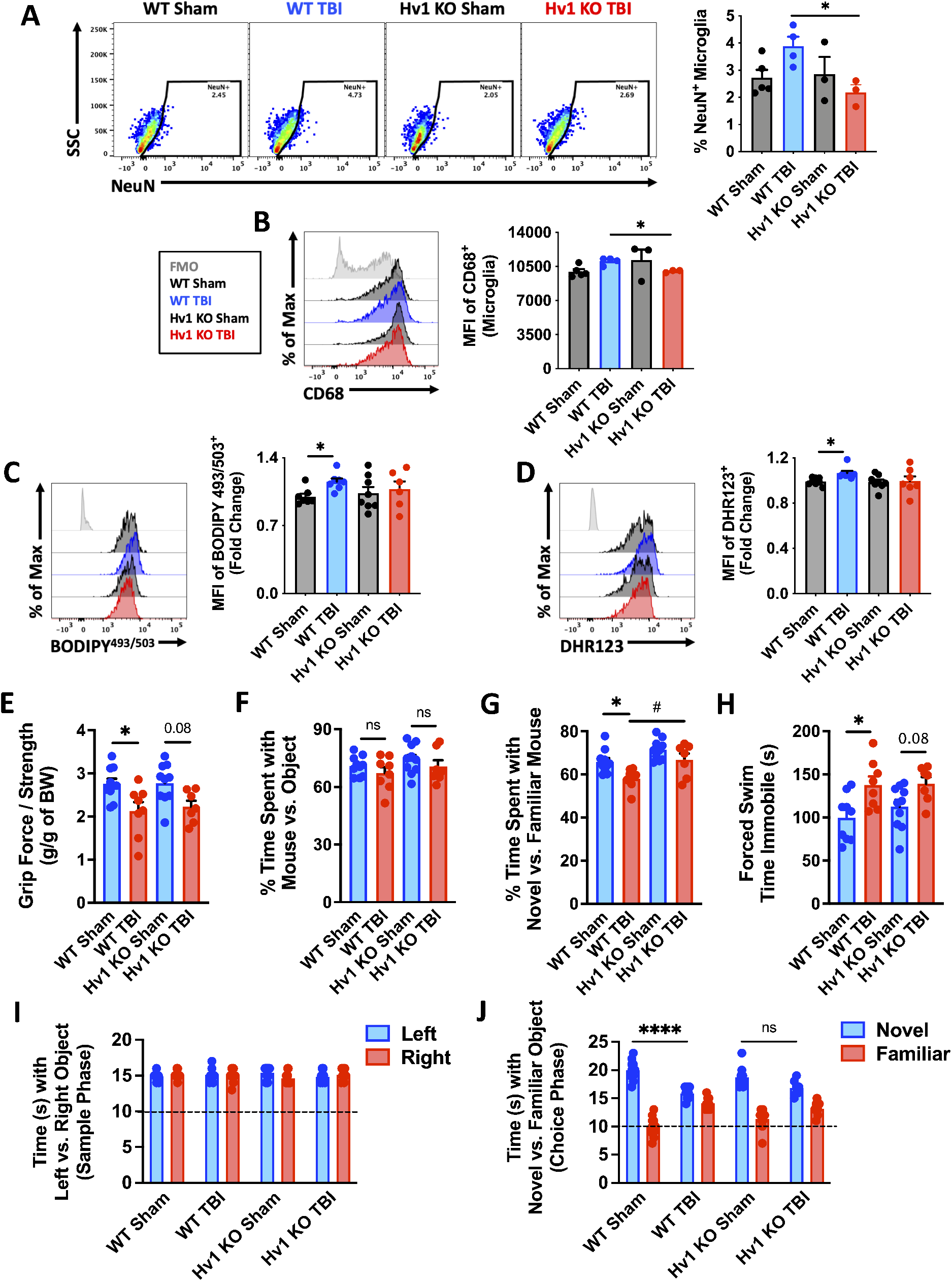
Hv1-driven oxidative stress promotes induction of the AF-related microglia phenotype and worsens long-term functional recovery after TBI. Injured Hv1^-/-^ (KO) and wildtype (WT) littermate control mice were behaviorally evaluated at 11 months post-TBI and microglia were immunophenotyped at 12 months. (**A**) A representative dot plot of microglia containing NeuN antigen at one year post-TBI in WT and Hv1 KO mice. Quantification of phagocytic microglia is shown (right). **(B)** A representative histogram shows the relative expression level of the phagocytosis marker CD68 in microglia. The MFI of CD68-positive microglia is quantified (right). **(C)** A representative histogram shows the relative level of neutral lipids in in microglia at one year after TBI. The MFI of BODIPY-positive microglia is quantified (right). **(D)** A representative histogram shows the relative level reactive oxygen species production in microglia using the DHR123 fluorescent probe. MFI quantification of DHR123-positive microglia is shown (right). **(E)** Forelimb grip strength was significantly reduced in WT but not Hv1 KO mice after TBI. **(F)** The percent of time spent interacting with a mouse versus object in the social recognition test. **(G)** The percent of time spent interacting with a novel stranger mouse versus familiar mouse in the social recognition test. **(H)** Time spent immobile during the forced swim test is shown for all groups. **(I)** The time spent interacting with left versus right objects during the sample phase of the object recognition test. **(J)** The time spent interacting with a novel versus familiar object during the choice phase of the object recognition test. For all behavioral experiments, N=7-10/group. For data (**A-B**), N=3-5/group. Data for (**C-D**) are representative of two independent experiments combined and expressed as a fold change to WT Sham values. For all histograms (**B-D**), FMO controls are shown in light gray, while all other groups are shown with bold outline as color coded according to the order of the bar graph axis labels (also shown in the figure legend). A vertical fiducial line is included for reference. Abbreviations: BW body weight, g grams, KO knockout, FMO fluorescent minus one, ns not significant, Max Maximum, MFI mean fluorescence intensity, s seconds, SSC side scatter, TBI traumatic brain injury, WT wildtype. Data (**I-J**) were analyzed using two-way ANOVA with Bonferroni’s post-hoc correction. Data **(A-D, E-H)** were analyzed using one-way ANOVA with Bonferroni post-hoc correction for multiple comparisons (*p<0.01 and **p<0.001).

## Discussion

Here we present the first detailed immunophenotypic analysis of naturally occurring, age-associated autofluorescent (AF) microglia in brain. This subpopulation of microglia in older mice adopt a unique neurotoxic phenotype defined by increases in phagocytosis, oxidative stress levels, lysosomal content and autophagy markers, lipid and iron accumulation, metabolic alterations, pro-inflammatory cytokine production, and senescent-like features. These age-associated changes in AF microglia were not regulated at the transcriptional level, but were more pronounced under pathological conditions (e.g., h*APOE4*), and were reversed with pharmacologically-mediated cell replacement/turnover. Furthermore, we demonstrated that TBI accelerates the age of onset and tissue-wide distribution of this AF phenotype via a phagocyte-driven oxidative stress mechanism.

Our immunophenotypic characterization of age-related AF microglia converge on the recently described lipid droplet-accumulating microglia, or LDAM, which increase with age, have a pro-inflammatory phenotype, and are also associated with neurodegeneration (*26*). However, because the cellular organelles known as lipid droplets, which are involved in energy storage, were identified based on neutral lipid staining, it was notable that no link was made between LDAM and lipofuscin-associated autofluorescence, which also contains neutral lipids, localize in and around lysosomes, and increases with age and neurodegeneration. In a related study, Burns et al. demonstrated that the AF signal in microglia increases linearly with age and is commensurate with the size and complexity of lamp1-positive lysosomal storage bodies (*27*). Proteomic analysis of AF-positive microglia showed enrichment endolysosomal, autophagic and metabolic pathways, including upregulation of lysosomal degradation enzymes and proteins involved in phagosome maturation. Interestingly, the authors found that myelin, Fc receptor-mediated, and TREM2-mediated phagocytosis were not dominant mechanisms contributing to AF accumulation. Rather, they found that genetic disruption of lysosomal function accelerated the accumulation of storage bodies in AF^+^ cells and led to impaired microglia physiology and cell death. Although there is strong support for the role of lysosomal enzymes in the degradation of ingested material in neurodegenerative diseases (e.g., lysosomal storages disorders), converging transcriptomic and proteomic data also suggest that AF microglia have functional differences in phagocytic capacity highlighted by an upregulation of biological pathways involved in the endocytosis process. However, few studies to date have demonstrated an increase in microglial phagocytosis in normal aged mice, despite numerous reports of increased CD68 expression, elevated NOX2 activity, DAM-associated gene signatures, and accompanying synaptic loss and myelin thinning (*28*). Our discovery that age-associated AF microglia exhibit a hyperphagocytic phenotype is consistent with these features and provides a mechanistic convergence between aging and post-traumatic neurodegenerative disease.

Lipofuscin (LF) accumulates in post-mitotic cells at a rate which occurs inversely to longevity (*17*). Lipid-laden neural cells can be visualized adjacent to blood vessels or deeper in the brain cortical and striatal parenchyma of aging mice and have distinct phenotypes associated with inflammation and senescence (*29*). Because laboratory mice live a relatively sanitary life devoid of the pathological hallmarks associated with neurodegenerative disease in humans, it was thought LF accumulation was just a consequence of time (*30*). However, evidence that microglia within amyloid plaques in humans show increased LF deposition implies that this feature is pathological (*31*). In contrast, proliferative cells have been shown to efficiently dilute LF aggregates during cell division, showing low or no accumulation of the pigment (*7*). Interestingly, the accumulation of LF (i.e., AF phenotype) in microglia may then be indicative of replicative senescence, which may occur naturally or following waves of injury-induced proliferation. It is reasonable to consider that microglial turnover or proliferation in aged mice may effectively dilute LF within the brain or cell population, respectively. Indeed, a reversal of age-related cognitive and synaptic deficits in old mice following microglia elimination and repopulation (i.e., replacement/turnover) can be achieved using short-term treatment with the CSF1 inhibitor, PLX5622 (*22*). And whereas microglial turnover has been demonstrated to reduce AF, partially reverse the aging microglia transcriptome (*18*), and improve amyloid pathology (*32*), several questions remain as to the linkage of these changes and which specific populations, if any, are responsible for promoting these age-related neuropathologies. Our findings suggest that AF microglia are dysfunctional, neurotoxic cells that display a senescent-like phenotype. Notably, pharmacological depletion of microglia in aged mice effectively eliminated the AF population, partially restoring cellular and neurological functions.

The present study also demonstrated the close association of AF microglia with neuroinflammation, neurodegeneration, and neurological decline in aged mice. To date, there is no method available to selectively deplete LF/AF microglia that would support causality. However, our finding that AF microglia have a neurotoxic phenotype that can promote cytokine-mediated tissue inflammation, leukocyte recruitment, hippocampal neuron loss, and functional decline provides a plausible explanation for age-related inflammatory-driven neuropathologies in animals. Moreover, because falls at ground level are the leading cause of TBI in older individuals (*33*), and forced microglial turnover ameliorated gait deficits, the presence or abundance of AF microglia may not just worsen outcome, but also increase risk.

Our finding that moderate-to-severe TBI at young maturity or old age can promote and intensify the emergence of microglial AF is consistent with the notion of accelerated immune aging and suggests an inducible and shared phenotype that exists along a continuum of neurodegenerative disease risk or severity. Sustained increases in expression of the phagosome marker CD68 and positive regulator of phagocytosis, NOX2, have previously been shown in chronically injured mice as late as one year after TBI (*34*). Moreover, microglial phagocytosis of apoptotic neurons and synapses was evident for up to eight months after TBI, coincident with decreased synaptic protein expression in neurons (*35*). An age-related increase in chronic engulfment of neuronal and myelin antigens late after TBI has also been demonstrated and such changes were accompanied by an age-related increase in the tissue expression of DAM genes, microglial ROS production, white matter loss, and hippocampal neuron death (*36*). Thus, the AF phenotype may identify a subset of dysfunctional or neurotoxic microglia responsible for these molecular signatures within heterogenous cell and tissue samples. A direct comparison between AF microglia from normal aged, chronic TBI, and other neurodegenerative disease states is required to better understand the homology of phenotypic features and inflammatory signatures. However, because such studies may be fraught with confounding factors of age, genotype, comorbities, and disease staging, careful consideration is needed.

The observation that oxidative stress mediates autophagic dysregulation and formation of LF with advanced age is rationale and supported in the literature. ROS have been demonstrated to contribute to LF buildup and pro-inflammatory activation in neural cells, peripheral macrophages and microglia (*5, 17, 37-40*). That a similar phenomenon occurs after brain injury is not well documented, despite evidence for persistent basal elevation of NOX2, ROS production, oxidative stress, autophagic dysregulation, loss of proteostasis, and pro-inflammatory phenotype (*41–44*). NOX2-deficient mice exhibit acute neuroprotection and fewer neurological deficits after TBI (*45, 46*). Experimental manipulation of ROS production, the oxidative stress response, and autophagy have likewise been demonstrated to attenuate neuroinflammation, neuronal loss, and functional impairment. AF^hi^ microglia produced higher levels of NOX2 and ROS, but whether it was the cause or consequence of LF was not clear. Utilizing mice deficient in Hv1, a proton channel present on phagosomes that facilitates sustained NOX2-mediated ROS production in phagocytes, we had previously showed that neuroinflammation and neurodegeneration were significantly reduced after TBI (*47*). Here we show that Hv1 chronically drives oxidative stress in phagocytes, promotes engulfment of neuronal debris, increases LF, and worsens neurological outcome. It is possible that APOE4 genotype and oxidative stress work synergistically to augment the age-related injury AF phenotype, but further investigation is required.

In summary, our data suggest that AF reflects a pathological state in aging microglia associated with hyperphagocytosis and inflammatory neurodegeneration that can be further accelerated by TBI. This newly immunophenotypically defined AF microglial population may serve as a useful imaging biomarker of DAM/senescence that may be otherwise underrepresented or difficult to detect in whole tissue or bulk cell molecular analyses given its largely post-transcriptional/translational underpinnings. Mechanistically, *APOE4* genotype at old age and chronic injury-induced oxidative stress promote the AF phenotype. Finally, pharmacological intervention can eliminate the AF microglial signature, reduce inflammatory burden, improve neurological function, and prophylactically protect the brain against severe head impact.

## Materials and Methods

### Animals

Young adult (3-month-old) and aged (18-month-old) male C57BL/6 mice bred in-house from Charles River Laboratories were housed on sawdust bedding in a specific pathogen free facility (12 hours light/dark cycle). B6(SJL)-Apoe^tm1.1(APOE*4)Adiuj^/J (Stock No: 027894), homozygous for a human Apoe4 gene targeted replacement of the endogenous mouse Apoe gene (26), were obtained from the Jackson Laboratories. Homozygous Hv1 mice breeders were obtained from Dr. Long-Jun Wu’s laboratory at Mayo Clinic, Rochester, MN and maintained in the UMB animal facility. SLICK-A transgenic mice (Jackson Laboratories, B6.Cg-Tg(Thy1-cre/ERT2,-EYFP)AGfng/J; Stock No: 007606) were used for phagocytosis studies as previously described (*35*). All animals had access to chow and water *ad libitium*. Animal procedures were performed in accordance with NIH guidelines for the care and use of laboratory animals and approved by the Animal Care Committee of the University of Maryland School of Medicine.

### PLX5622 administration

PLX5622 was provided by Plexxikon and formulated in AIN-76A rodent chow by Research Diets at a concentration of 1200 ppm (*48, 49*). Standard AIN-76A diet was provided as Vehicle (Veh) control. Mice were provided *ad libitum* access to PLX5622 or Veh diet for 3 weeks to deplete microglia. After 3 weeks of depletion, mice were placed back on normal chow for either 4 or 12 weeks to allow for repopulation. This dose and time resulted in depletion of 95% of microglia and significant depletion of CD115/CSF1R-expressing myeloid cells in other peripheral organs consistent with previous studies (**Supplemental Fig. 1**) (*50*).

### Controlled cortical impact

Our custom-designed controlled cortical impact (CCI) injury device consists of a microprocessor-controlled pneumatic impactor with a 3.5 mm diameter tip as previously described (*49*). Briefly, mice were anesthetized with isoflurane evaporated in a gas mixture containing 70% N_2_O and 30% O_2_ and administered through a nose mask. Mice were placed on a heated pad and core body temperature was maintained at 37°C. The head was mounted in a stereotaxic frame, a 10 mm midline incision was made over the skull and the skin and fascia were reflected. A 5 mm craniotomy was made on the central aspect of the left parietal bone. The impounder tip of the injury device was then extended to its full stroke distance (44 mm), positioned to the surface of the exposed dura, and reset to impact the cortical surface. Moderate-level CCI was induced using an impactor velocity of 6 m/s, deformation depth of 1 mm, and a dwell time of 50 ms (*49*). After injury, the incision was closed with interrupted 6-0 silk sutures, anesthesia was terminated, and the animal was placed into a heated cage to maintain normal core temperature for 45 minutes post-injury. Sham animals underwent the same procedure as CCI mice except for craniotomy and cortical impact. In the 14d and Hv1 TBI cohorts, an injury of moderate severity was induced by a TBI-0310 Head Impactor (Precision Systems and Instrumentation) with a 3.5-mm diameter tip followed by impact velocity of 4.3 m/s and a displacement depth of 1.2 mm (*36, 47*).

### Flow cytometry and *ex vivo* functional assays

Following transcardial perfusion with 40 mL of ice-cold sterile PBS, the intact brain was harvested, and the ipsilateral hemisphere isolated by removing the olfactory bulbs and cerebellum. Brain hemispheres were mechanically digested using a razor blade to mince tissue and were passed through a 70-μm-filter using RPMI. CNS tissue was then enzymatically digested using DNAse (10 mg/mL; Roche), Collagenase/Dispase (1 mg/mL; Roche), and Papain (25 U; Worthington Biochemical) for 1 hour at 37°C in a shaking CO_2_ incubator (200 rpm). Tissue homogenates were centrifuged at 1500 rpm for 5 minutes at 4°C. The supernatant was discarded and the cells were resuspended in 70% Percoll™ (GE Healthcare) and underlayed in 30% Percoll™. This gradient was centrifuged at 500 g for 20 minutes at 21 °C. Myelin was removed by suction and cells at the interface were collected. Leukocytes were washed and blocked with mouse Fc Block (eBioscience, clone 93) prior to staining with primary antibody-conjugated flourophores (CD45-eF450 (30-F11), CD11b-APCeF780 (M1/70), Ly6C-APC (HK1.4), and Ly6G-PE (1A8) were purchased from eBioscience, whereas CD45-PerCP-Cy5.5 (30-F11) and CD11b-PerCP-Cy5.5 (M1/70) were purchased from Biolegend. For live/dead cell discrimination, a fixable viability dye, Zombie Aqua™ (Biolegend), was dissolved in DMSO according to the manufacturer’s instructions and added to cells in a final concentration of 1:50. Data were acquired on a LSRII using FACsDiva 6.0 (BD Biosciences) and analyzed using FlowJo (Tree Star). A standardized gating strategy was used to identify microglia (CD45^int^CD11b^+^Ly6C^−^) and brain-infiltrating myeloid cells (CD45^hi^CD11b^+^), including monocyte (CD45^hi^CD11b^+^Ly6C^+^Ly6G^−^) and neutrophil (CD45^hi^CD11b^+^Ly6C^+^Ly6G^+^) populations as previously described (*51*). Cells were also stained with the surface markers CD115 (CSF1R)-AF488, BioLegend (AFS98); CD16/32-APC, BioLegend (93); and CD369 (CLEC7A)-APC, Biolegend (RH1). Cell-specific fluorescence minus one (FMO) controls were used to determine the positivity of each antibody. Cell count estimations were performed using CountBright™ absolute counting beads (20 μL/test; Invitrogen, Carlsbad, CA, USA) according to the manufacturer’s instructions.

Intracellular staining was performed using a fixation/permeabilization kit (BD Biosciences) and the following antibodies: p16^ink4a^-APC, StressMarq Biosciences (SPC-1280D); NOX2-AF647, Bioss Antibodies (bs-3889R); GAPDH-AF488, BioLegend (W17079A); Lamp1-PerCPCy5.5, BioLegend (1D4B); Lamp2-PE, BioLegend (M3/84); NeuN-PE, Millipore Sigma (A60); MAP2-AF594, BioLegend (SMI 52); Myelin CNPase-AF647, BioLegend (SMI 91); and CD68-PE-Cy7, BioLegend (FA-11).

For intracellular cytokine staining, leukocytes were collected as described above, and 1 μL of GolgiPlug containing brefeldin A (BD Biosciences) was added to 500 μL complete RPMI. Cells were then resuspended in Fc Block, stained for surface antigens and washed in 100 μL of fixation/permeabilization solution (BD Biosciences) for 20 minutes. Cells were washed twice in 500 μL Permeabilization/Wash buffer (BD Biosciences) and resuspended in an intracellular antibody cocktail containing cytokine antibodies (MMP-9-R-PE, StressMarq Biosciences (SMC-396D); TNF-PE-Cy7, eBioscience (MP6-XT22); IL-1α-PE, BioLegend (ALF-161); and IL-1β-PerCP-eF710, eBioscience (NJTEN3) and fixed.

For ROS detection, leukocytes were incubated with dihydrorhodamine (DHR) 123 (5mM, Life Technologies/Invitrogen) or H_2_DCFDA (DCF, 5 μM; ThermoFisher Scientific), cell-permeable fluorogenic probes. Cells were loaded for 20 minutes at 37°C, washed three times with FACS buffer (without NaAz), and then stained for surface markers including viability stain. Lipid peroxidation was measured using BODIPY 581/591 C11^+^ (Invitrogen). LC3-associated autophagic vesicles were measured using the CYTO-ID Autophagy Detection Kit (Enzo Life Sciences) according to the manufacturer’s instructions. Lysosomes were labeled using LysoTracker Green DND-26 (Invitrogen). Intracellular myelin content was stained with FluoroMyelin Red (Invitrogen). Neutral lipids and lipid droplets were measured using BODIPY 493/503 (Invitrogen), Lipi-Blue (Dojindo, LD01-10). Intracellular iron levels were measured using FerroOrange (Dojindo). Relative intracellular pH was measured using Protonex Red 600 (AAT Bioquest, 5 μM). Mitochondrial membrane potential was measured using MitoSpy Red dye (BioLegend) according to the company’s protocol. Glucose uptake was measured using 2NBD-glucose uptake cell-based assay kit (Cayman Chemical).

Microglial phagocytosis of apoptotic neurons was performed as described by Ritzel et al. (*35*). Briefly, YFP-positive neurons were isolated from the cortices of an adult SLICK transgenic mouse and then exposed to heat shock for 5 minutes at 60 °C, washed in HBSS, resuspended in 5 mL of RPMI, and kept on ice. Soon after, 50 μL of feeder YFP-positive cells were incubated with freshly isolated microglia for 45 minutes at 37 °C. Afterward, the cells were washed three times with 1 mL PBS, stained and fixed as above, and collected on the cytometer.

### FACS sorting of microglia

Living CD45^+^CD11b^+^Ly6C^-^ Microglia were sorted (100-μm nozzle/18 psi) using a FACS Aria II (BD) by the University of Maryland Greenebaum Comprehensive Cancer Center Flow Cytometry Shared Service Core. Prior to sorting, brain hemispheres were processed as above using FACS buffer without sodium azide or fixation. Brain leukocytes were washed and blocked with mouse Fc Block (ThermoFisher, clone 93) prior to staining with primary antibody-conjugated flourophores. CD45-eF450 (30-F11), CD11b-APCeF780 (M1/70), and Ly6C-APC (HK1.4) were purchased from ThermoFisher. For live/dead cell discrimination, the viability dye, 7-AAD (Biolegend), was diluted at 5uL in 100uL sample. Data were acquired and microglia were FAC-sorted on a FACS ARIA II using FACsDiva 6.0 (BD Biosciences). Purified microglia from individual hemispheres were collected in 300 μl of RNAprotect cell reagent (Qiagen) with β-mercaptoethanol (Sigma) into a standard sterile 1.5 mL RNAse-free Eppendorf microcentrifuge tube, vortexed for 30 s, snap frozen in dry ice, and then stored in −80 °C until RNA extraction. Each tube was later thawed, vortexed again, and poured over a column using the RNA kit as described. All microglia were sorted on two separate days within the same week using the same materials and reagents. On the first sort day young bulk (i.e., AF^lo^), old AF^lo^, and old AF^hi^ microglia populations were collected (N=4/group). Microglial AF levels were quantified for each FACS-sorted group (**Supplemental Fig. 2**). On the second day, young + vehicle, old + Veh, and old + PLX5622 (i.e., 3 weeks elimination + 4 weeks repopulation) bulk microglia populations were collected (N=4/group). Approximately 5,000 to 55,000 microglia were collected for each tube per group.

### RNA sequencing and transcriptomic analysis

FACS sorted microglia cells were flash frozen in dry ice into 300 μL RNAprotect cell reagent (Qiagen). Total RNA was extracted using RNeasy mini kit (Qiagen). RNA samples were sent to the Institute for Genome Sciences at University of Maryland School of Medicine for RNA quality test and RNA sequencing. The RNA quality was analyzed using low-input Pico chip on Agilent 2100 Bioanalyzer. Amplification of cDNA from the total RNA samples were performed using Ovation RNA-seq system kit (Tecan), following by low input RNA-seq library preparation. RNA sequencing of all libraries was performed on Illumina HiSeq4000. The data were converted to FastQ format.

Bioinformatics analyses were performed on the High Performance Computing Facility taki CPU cluster at University of Maryland Baltimore County. Data quality were analyzed using FastQC v0.11.8 (*Available online at:* http://www.bioinformatics.babraham.ac.uk/projects/fastqc/). Ribosomal RNA were filtered using SortMeRNA v2.1b (*52*). Transcript-level abundance was quantified using the quasi-mapping-based mode in Salmon v1.0.0 (*53*), mapping to GRCm38 (mm10) mouse reference genome. Transcript-level abundance were aggregated to gene-level abundance using tximport (*54*). Genes with abundance of 0 in any samples were filtered out from further analysis. DESeq2 (*55*) was used for differential expression analyses. P-values were adjusted using the Holm-Bonferroni method and genes with adjusted p-value < 0.1 were defined as significant differentially expressed genes. Principal component analysis was performed using the mixOmics package (*56*). Tximport, DESeq2, and mixOmics packages were run in R v3.6.1 in RStudio v1.2. Sample-level enrichment analysis (SLEA) was performed as previously described (*57*). Briefly, the mean expression of 10,000 random sets of genes the same size as the scored gene set were calculated as used as the null distribution for each sample. The mean expression of the scored gene set was then calculated and transformed to z-score based on the null distribution (SLEA score). Gene sets were used from the Gene Ontology Biological Process database for phagocytosis (M16307), regulation of autophagy (M10281), and regulation of reactive oxygen species biosynthetic process (M15379). DAM/senescence gene set was used from Hu et al. 2021 (*58*).

### Neurobehavioral testing

The following behavior tests were performed with mice and group information blinded to the operators. To minimize stress and fatigue, each test was performed on a different day.

#### Motor function

##### Rotarod

Locomotor function and coordination were assessed using a rotarod as previously described (*59*). The mouse was placed on a rotarod device (IITC Life Science, Inc.), and their latency to falling off the accelerating rotarod was recorded. The acceleration settings for the device were 4 to 40 rpm over 90 seconds, with each trial lasting for a maximum of 300 seconds. Individual scores from three trials were averaged and evaluated relative to their baseline latencies.

##### Grip strength

Grip strength was measured using a digital grip strength meter (Bioseb BP, In Vivo Research Instruments) as previously described (*36, 60*). Forelimb grip strength was measured from the mouse using both the ipsilateral and contralateral forepaws together. The mouse was held by its tail, the forelimbs were placed on the grasping metal wire grid and the mouse gripped the wire grid attached to the force transducer. Once the grip was secured, the animal was slowly pulled away from the bar. The maximal average force exerted on the grip strength meter by both forepaws was averaged from 10 trials per day for each mouse.

##### Gait dynamics

Analysis of gait and posture was performed with the CatWalk XT automated system as mentioned in our previous publications (Noldus) (*35, 61*). Acquisition of data took place in a darkened room with red light. The Catwalk apparatus records print position, gait postures and weight distribution through its green illuminated walkway. A minimum of 3 valid runs, complete crossings with no turns or scaling of sidewalls, were obtained for each tested mouse. Runs that didn’t comply to the preset standards were excluded from the final analysis. The Regularity Index (%) expresses the number of normal step sequence patterns relative to the total number of paw placements. The Regularity Index is a fractional measure of inter-paw coordination. In healthy, fully coordinated animals its value is 100%. Stand (s) is the duration of contact with the glass plate of the print. Stride Length (cm) is the distance (in Distance Units) between successive placements of the same paw. Swing Speed (cm/s) is the speed (Distance Unit/second) of the paw during Swing (Swing Speed = Stride Length / Swing). The Body Speed (cm/s) of a step cycle of a specific paw is calculated by dividing the distance that the animal’s body traveled from one initial contact of that paw to the next by the time to travel that distance. Step Cycle (s) is the time in seconds between two consecutive Initial Contacts of the same paw (Step Cycle = Stand + Swing).

#### Cognitive function

##### Novel object recognition (NOR) task

For testing non-hippocampal mediated memory, mice in Cohort 3 underwent NOR as previously described (*36, 62*). Mice were tested in an open field apparatus after a 5 minute habituation period on the first day. The time spent with two identical objects was recorded using ANY-maze software (Stoelting) on the second day of testing, and one of the familiar objects were switched out with a novel object on the third day. Testing stopped after each mouse went through a sum total of 30 s exploration time. Since mice would inherently prefer to explore novel objects, a preference for the novel object with an exploration time of more than 15 s was considered as having intact learning and memory skills.

##### Y-maze test

This task was used to assess the hippocampus-dependent spatial working memory of mice as previously described (*47*). The Y-maze (Stoelting, Wood Dale, IL) consisted of three identical arms, each arm 35 cm long, 5 cm wide, and 10 cm high, at an angle of 120° with respect to the other arms. One arm was randomly selected as the “start” arm, and the mouse was placed within and allowed to explore the maze freely for 5 minutes. Arm entries (arms A–C) were recorded by analyzing mouse activity using ANY-maze software (Stoelting). An arm entry was attributed when all four paws of the mouse entered the arm, and an alternation was designated when the mouse entered three different arms consecutively. The percentage of alternation was calculated as follows: total alternations x 100/(total arm entries - 2). If a mouse scored significantly greater than 50% alternations (the chance level for choosing the unfamiliar arm), this was indicative of spatial working memory.

#### Depressive-like behavior

##### Social recognition (SR) task

Social recognition test was performed for assessment of sociability function as previously described (*35, 47*). Using a three-chambered rectangular apparatus made of Plexiglas with each one at equal size (20 x 40 x 23 cm). An opening between the walls allows for free access to each chamber, which contains two identical wire mesh cup containers. Before testing, each mouse was single housed overnight. On the first day, the tested mice were placed in the apparatus with two empty cups for a 10 minute habituation period. On the second day, a stranger mouse was introduced and randomly placed inside one of the empty cups in either the left- or right-side chamber while the other cup was left empty. The tested mouse started from the middle chamber and allowed to freely explore all three chambers for an exploration period of 10 minutes.

Afterwards, a second unfamiliar, stranger was placed inside the previously empty cup. The test subject was once again allowed to freely explore all three chambers for a period of 10 minutes. Exploration time that the subject mice spent with each cup versus stranger mouse was recorded using ANY-maze software (Stoelting). Since a socially functional mouse would naturally seek out unfamiliar mice for interaction, the test subject was considered capable of social recognition if index for novel mouse scored higher than 50%.

##### Tail suspension (TS) test

The TS test assesses depression-like behavior in mice and is based on the observation that mice develop an immobile posture when placed in an inescapable hemodynamic stress of being hung by their tail. The TS was performed as previously described (*63, 64*). Each mouse in Cohort 3 was suspended at a height of 28 cm using 3M adhesive tape. The tip of the mouse tail was not wrapped around the rod while being suspended. The duration of immobility was recorded throughout the 5-minute test period. The definition of immobility is passive hanging and complete motionlessness. Foam padding (3” deep) was placed under the beam in case animals fall from the beam during the experiment.

##### Forced swim (FS) test

FS testing is one of the most commonly used assays for the study of depressive-like behavior in rodents, and was performed as previously described (*35, 47*). Mice were placed in transparent plastic cylinder (45 cm high X 20 cm diameter) filled with water (23 ± 2 °C; 28 cm in depth) for 6 minutes. The duration of immobility was recorded using ANY-maze software (Stoelting).

### Multiplex ELISA

Cerebellar tissue and the cortex tissue surrounding the lesion area were excised. Tissues were loaded into tissue homogenizing tubes (Bertin Corp) on ice with 200 μL NP40 cell lysis buffer supplemented with 1x Halt™ protease and phosphatase inhibitor cocktail (ThermoFisher Scientific), and then homogenized using a Fastprep FP120 homogenizer (Thermo Savant Bio 101) at 6.5 m/s in a 4 °C cold room for 2 minutes, with 30 seconds incubation on ice at 30 second intervals. Lysates were transferred into 1.5 mL centrifuge tubes and spun at 13,000 rpm for 10 minutes at 4 °C. Cleared supernatant was collected and protein concentration was quantified using Pierce BCA protein assay kit (ThermoFisher Scientific). Multiplex ELISA was performed on a Luminex FlexMAP 3D (Millipore Sigma) using a premixed 32-plex cytokine magnetic bead panel (Millipore Sigma) following the kit protocol. Undiluted cleared tissue lysate (25 μL) was assayed in technical duplicates. Cytokine concentrations were determined by interpolation to known standard curves followed by normalization to total protein concentration. Samples below the limit of detection were set to the lowest standard concentration. Analytes that were not within detectable limits or showed no change after injury are not shown.

### Lesion volume, neuronal counting, immunohistochemistry (IHC), and quantification

At 2 weeks post-injury, mice were perfused intracardially with normal saline followed by 4% paraformaldehyde solution. The brain was extracted and embedded in Tissue-Tek OCT compound (Sakura). Serial sections of 40μm thickness were placed on Superfrost Plus slides (ThermoFisher). Every eighth section was selected for analysis beginning from the foremost section. Lesion volume was performed after staining with cresyl violet (FD NeuroTechnologies). Quantification of the lesion volume was performed with the Stereoinvestigator software (MBF Biosciences) as previously described (*47*). By outlining the missing tissue on the injured hemisphere, the software was able to estimate lesion volume with the Cavalieri method at a grid spacing of 0.1mm. For quantifications of neuronal cell loss post injury, the optical fractionator method of unbiased stereology was used in the Stereoinvestigator software as previously described (*47, 64*). In the dentate gyrus region of the hippocampus, every fourth section was analyzed for a total of eight sections per mouse, beginning at bregma −2.15 mm and ending at −3.39 mm. The number of surviving neurons in each field was divided by the volume of the region of interest to obtain the neuronal cells density expressed in cells/mm^3^.

For IHC, brain slices were washed 3 times with phosphate-buffered saline (PBS) followed by blocking in 5 % normal goat serum containing 0.3% Triton X-100 in PBS for 2 hours. The primary antibodies were added into blocking buffer and incubated with brain sections overnight at 4 ℃. Sections were rinsed with PBS 3 times then incubated with secondary antibodies in blocking buffer for 2 hours at room temperature, followed by counterstaining with 4’,6-diamidino-2-phenylinodole (DAPI, Sigma-Aldrich) for 10 minutes. After washing with PBS, sections were mounted onto glass slides with coverslips using an anti-fade Hydromount solution (National Diagnostics). The following primary and secondary antibodies were used: rabbit anti-Iba1 (1:1000, Wako); rat anti-CD68 (1:1000, BioLegend); Alexa Fluor^TM^ 546 goat anti-rabbit IgG (1:800, Invitrogen); Alexa Fluor^TM^ 647 goat anti-rat IgG (1:800, Invitrogen). The green channel was left blank for imaging of AF.

Eight sections from each mouse in Veh/TBI and PLX/TBI groups were stained and imaged and a total of N=8 mice/group were used in the analysis. Images were acquired using a Leica TCS SP5 II Tunable Spectral Confocal microscope system (Leica Microsystems). For each section, four cortex regions in the ipsilateral side close to the lesion core were imaged as well as four regions in the contralateral cortex by identical imaging parameters with 3 times zooming in under a 20X lens. Using the NIH ImageJ software (1.53), a single view, an area of 258.84μm x 258.84 μm with 512 x 512 resolution, was formed by the Z-axis projection of 10 layers covering the entire depth. The number of Iba1^+^ cells, CD68^+^ counts, or autofluorescence particles, were quantified automatically by the “Analyze Particles” function in ImageJ after background subtraction and signal filtering. For each animal, the mean value of four regions on the same side was taken into statistical analysis. The signal colocalization of autofluorescence with Iba1 or CD68 was confirmed by the “Orthogonal View” function in ImageJ (*36, 64*). All IHC experiments were performed with appropriate positive control tissue, as well as primary/secondary only negative controls. IHC analysis was performed by an investigator blinded to experimental groups.

### Statistical analysis

Data from individual experiments are presented as mean ± S.E.M. One-way ANOVA was employed with Tukey’s multiple comparisons test, and group effects were determined by two-way ANOVA analysis with Sidak’s or Tukey’s post-hoc correction for multiple comparisons. Behavioral data were analyzed by two-way repeated measures ANOVA. All behavioral, histological, and *ex vivo* studies were performed by an investigator blinded to genotype, treatment, and surgical condition. Statistical analysis was performed using GraphPad Prism Software v. 9.0 (GraphPad Software, Inc., La Jolla, CA). p≤0.05 was considered statistically significant.

## Acknowledgements

This work was supported by grants from the National Institutes of Health [F32NS105355 (RMR), K99NS116032 (RMR), RF1NS110637 (JW), R01NS094527 (JW), R01NS110635 (AIF/JW), R01NS110825 (JW), RF1AG069196 (SL)], and Science Foundation Ireland (SFI 17/FRL/4860 to DJL). The authors thank Plexxikon Inc. for the use of PLX5622 and Dr. Long-Jun Wu at Mayo Clinic for the use of Hv1 KO mice. We thank Julie Faden, Victoria Meadows, Jordan Carter, Hui Li, and Wesley Shoap for their assistance with animal care and behavioral experiments. We would also like to thank Ethan Glaser, Nicholas Braganca, Nivedita Hegdekar, Niaz Khan, and Courtney Colson for their assistance with flow cytometry experiments. We appreciate the advice of Dr. Evan Jellison, Assistant Professor of Immunology and Director of the Flow Cytometry Core at UCONN Health, Farmington, Connecticut.

## Author contributions

RMR conceived this study, designed, and conducted most of the experiments, supervised the project, and wrote the manuscript. YL performed the behavioral experiments. RMR, YL, and SJD performed the flow cytometry experiments. ZL performed the IHC analysis. YL and RS performed the histological analysis. JH and RJH performed the animal surgeries. YJ and GS analyzed RNAseq data. YJ performed the multiplex ELISA. GS, SL, AIF, and BAS provided feedback on the manuscript. JW and DJL contributed to study conception and design, revised manuscript. All authors have critically read and commented on the final paper.

## Data and materials availability

All data needed to evaluate the conclusions in the paper are present in the paper and/or the Supplementary Materials. Additional data related to this paper may be requested from the corresponding authors. RNAseq data files will be deposited into the publicly available Gene Expression Omnibus (GEO), upon acceptance of manuscript.

## Ethics declarations

All animal procedures were conducted in accordance with the guidelines set forth by the National Institutes of Health and the University of Maryland, Baltimore Institutional Animal Care and Use Committee, who approved the study protocol.

