## Supplemental Figure 1 for "Brain injury accelerates the onset of a reversible age-related microglial phenotype associated with hyperphagocytosis and inflammatory neurodegeneration"

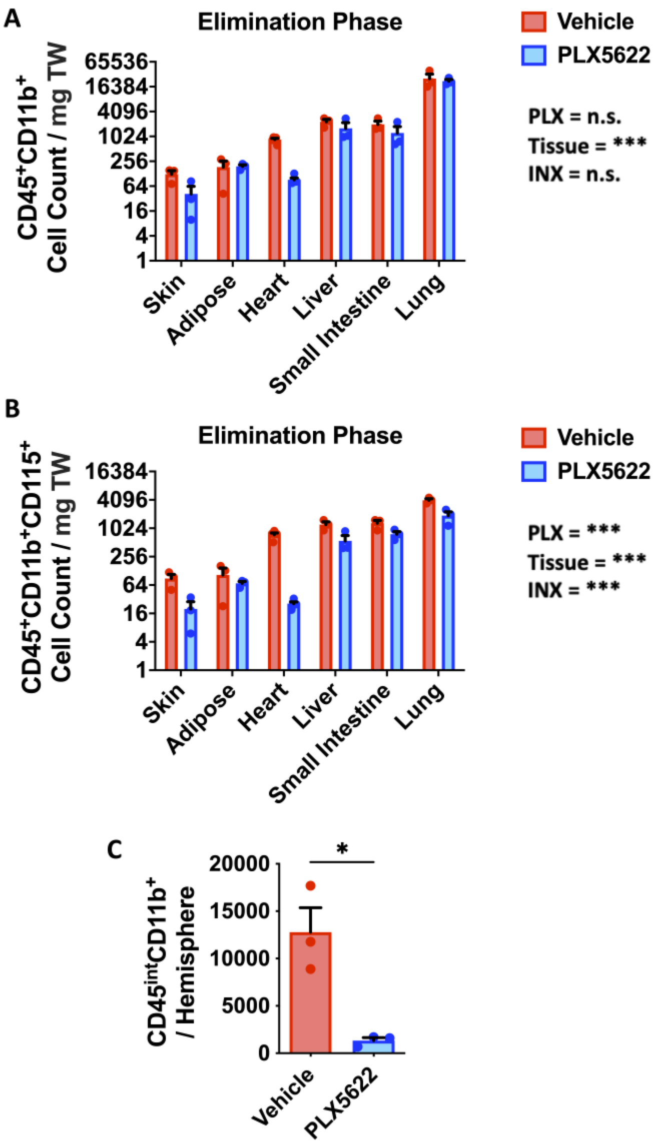


**Fig. S1. Peripheral immune composition following short-term CSF1R inhibition with PLX5622.**

**(A)** Cell counts of $CD45^{+}CD11b^{+}$ immune cells in multiple organs was quantified using flow cytometry at two-weeks after continuous treatment with PLX5622 in young male mice. **(B)** Further refinement of the gating strategy to include $CD115^{+}$ cells, which express CSF1R, show even higher significance as evidenced by two-way ANOVA group interaction. **(C)** The number of brain microglia was significantly reduced during PLX5622 treatment. Abbreviations: INX interaction, mg milligrams, PLX Plexxikon 5622, TW tissue weight. Data **(A-B)** were analyzed using two-way ANOVA with Bonferroni's post-hoc correction. Data **(C)** were analyzed using Student's t-test (*$p<0.01$ and ***$p<0.001$).
