## Supplemental Figure 2 for "Brain injury accelerates the onset of a reversible age-related microglial phenotype associated with hyperphagocytosis and inflammatory neurodegeneration"

**4wks post-Repopulation**

**Cellular Profile of Sequenced Microglia**

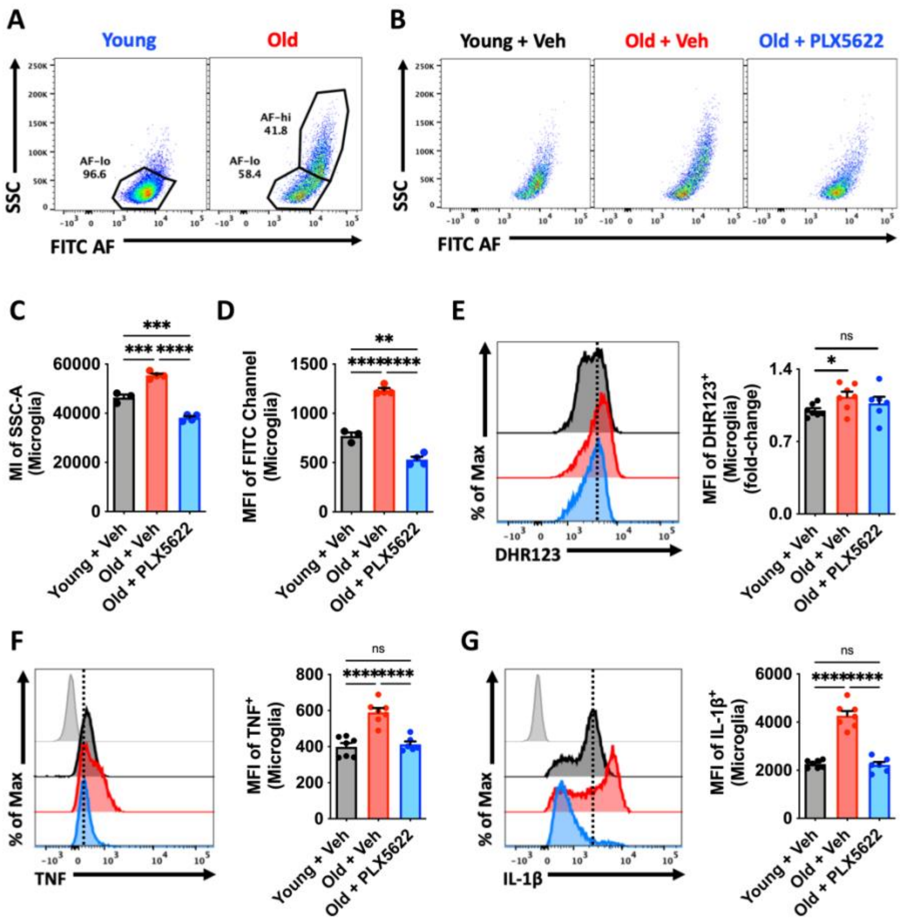


**Fig. S2. Cellular profile of microglia sorted for RNA sequencing.**

Two sets of experiments were performed to sort microglia for downstream RNA sequencing. In the first, living $CD45^{int}CD11b^{+}Ly6C^{-}$ microglia from young and old wildtype C57Bl/6 male mice were sorted based on the relative level of cellular autofluorescence in the FITC channel **(A)**. FACS sorted microglial subsets (e.g., Old $AF^{lo}$ and Old $AF^{hi}$) in the first experiment were confirmed autofluorescent based on relative intensity in the FITC channel with respect to the young (i.e., $AF^{lo}$) group. In the second experiment, rather than sorting by subset, bulk $CD45^{int}Cd11b^{+}Ly6C^{-}$ microglia from young and old vehicle groups and old treatment groups were sorted for comparison **(B)**. Validation of the PLX5622 treatment effect in the bulk sorted groups was performed to show the significant decrease in **(C)** cellular granularity and **(D)** autofluorescence level in microglia following four weeks of repopulation. Microglial production of reactive oxygen species **(E)** and the pro-inflammatory cytokines, TNF **(F)** and IL1β **(G)** are shown. Abbreviations: AF autofluorescent, hi high, lo low, MI mean intensity, MFI mean fluorescence intensity, SSC side scatter, Veh vehicle. Data **(C-G)** were analyzed using one-way ANOVA with multiple comparisons (**p<0.01, ***p<0.001 and ****p<0.0001).
