## Supplemental Figure 3 for "Brain injury accelerates the onset of a reversible age-related microglial phenotype associated with hyperphagocytosis and inflammatory neurodegeneration"

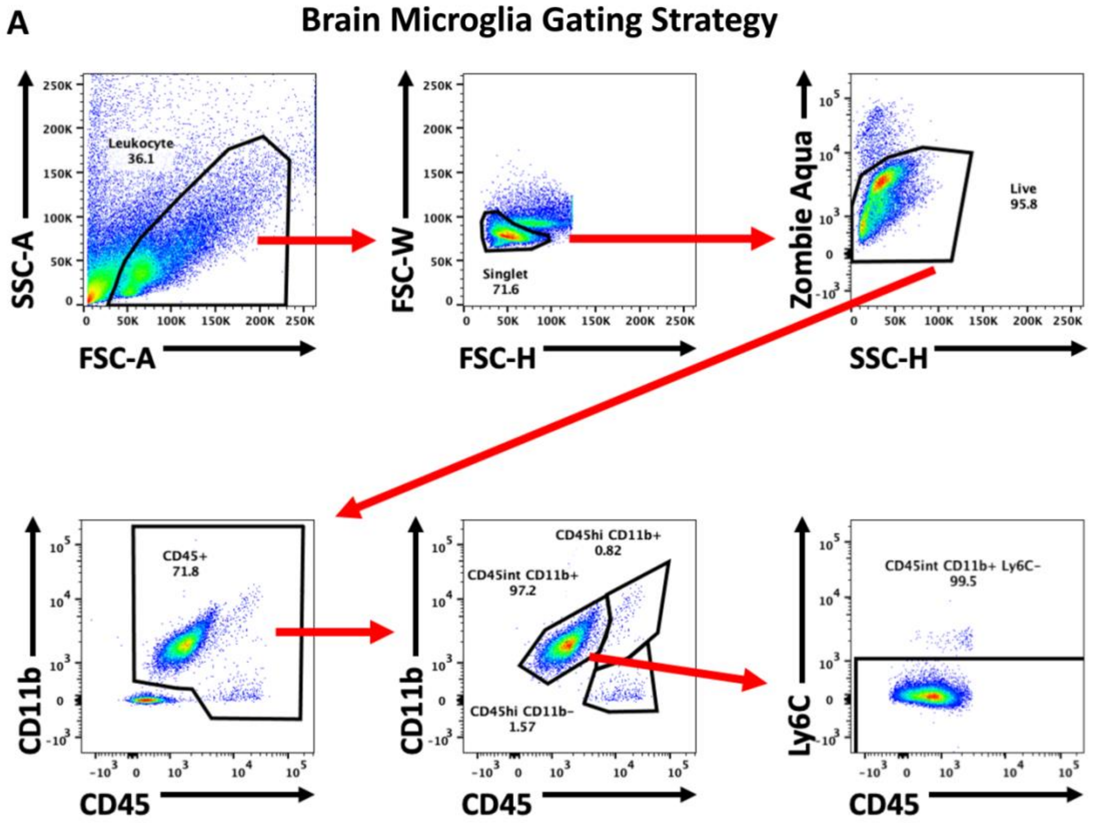


**Fig. S3. Flow cytometry identification of microglia.**

**(A)** Microglial cells in the brain were identified using the following gating strategy: leukocytes were first identified using a light scatter reference, then singlet gating, viable live cells, CD45 expression, $CD45^{int}CD11b^{+}$ expression, and finally, Ly6C-negative expression. Abbreviations: FSC forward scatter, hi high, int intermediated, SSC side scatter.
