## Supplemental Figure 4 for "Brain injury accelerates the onset of a reversible age-related microglial phenotype associated with hyperphagocytosis and inflammatory neurodegeneration"

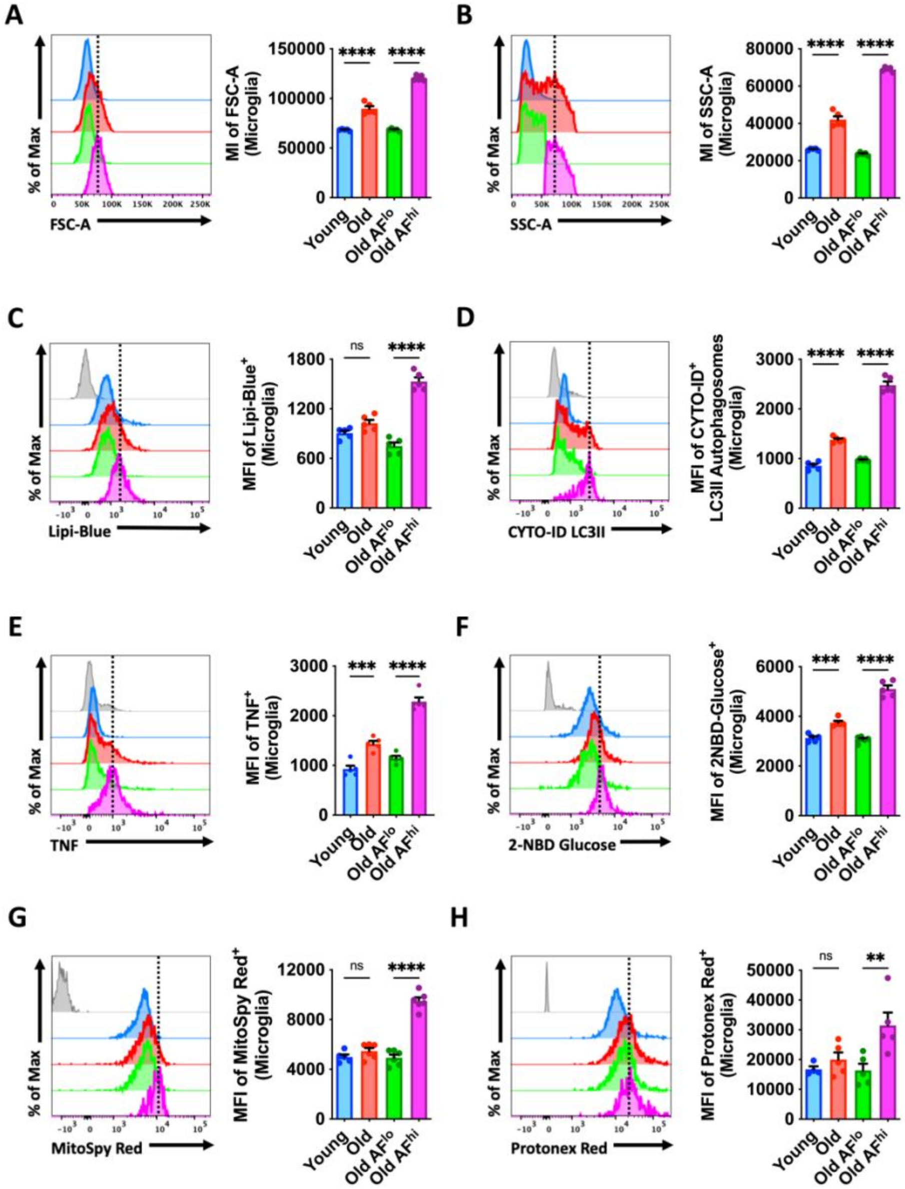


**Fig. S4. Functional evaluation of AF microglia from aged mice.**
Significant changes in cellular function of old AF$^{hi}$ versus old AF$^{lo}$ microglia subsets largely account for the age-related differences found in the general cell population. Representative histograms show that Old AF$^{hi}$ microglia exhibit increases in **(A)** cell volume, **(B)** cell granularity, (**C**) lipid content, **(D)** autophagosome formation, **(E)** TNF production, **(F)** glucose uptake, **(G)** mitochondrial membrane potential, and **(H)** cytosolic acidosis. Abbreviations: AF autofluorescent, FSC forward scatter, hi high, lo low, Max maximum, MFI mean fluorescence intensity, ns not significant, SSC side scatter. Data **(A-B)** were analyzed using one-way ANOVA with multiple comparisons (**p<0.01, ***p<0.001 and ****p<0.0001).
