## Supplemental Figure 5 for "Brain injury accelerates the onset of a reversible age-related microglial phenotype associated with hyperphagocytosis and inflammatory neurodegeneration"

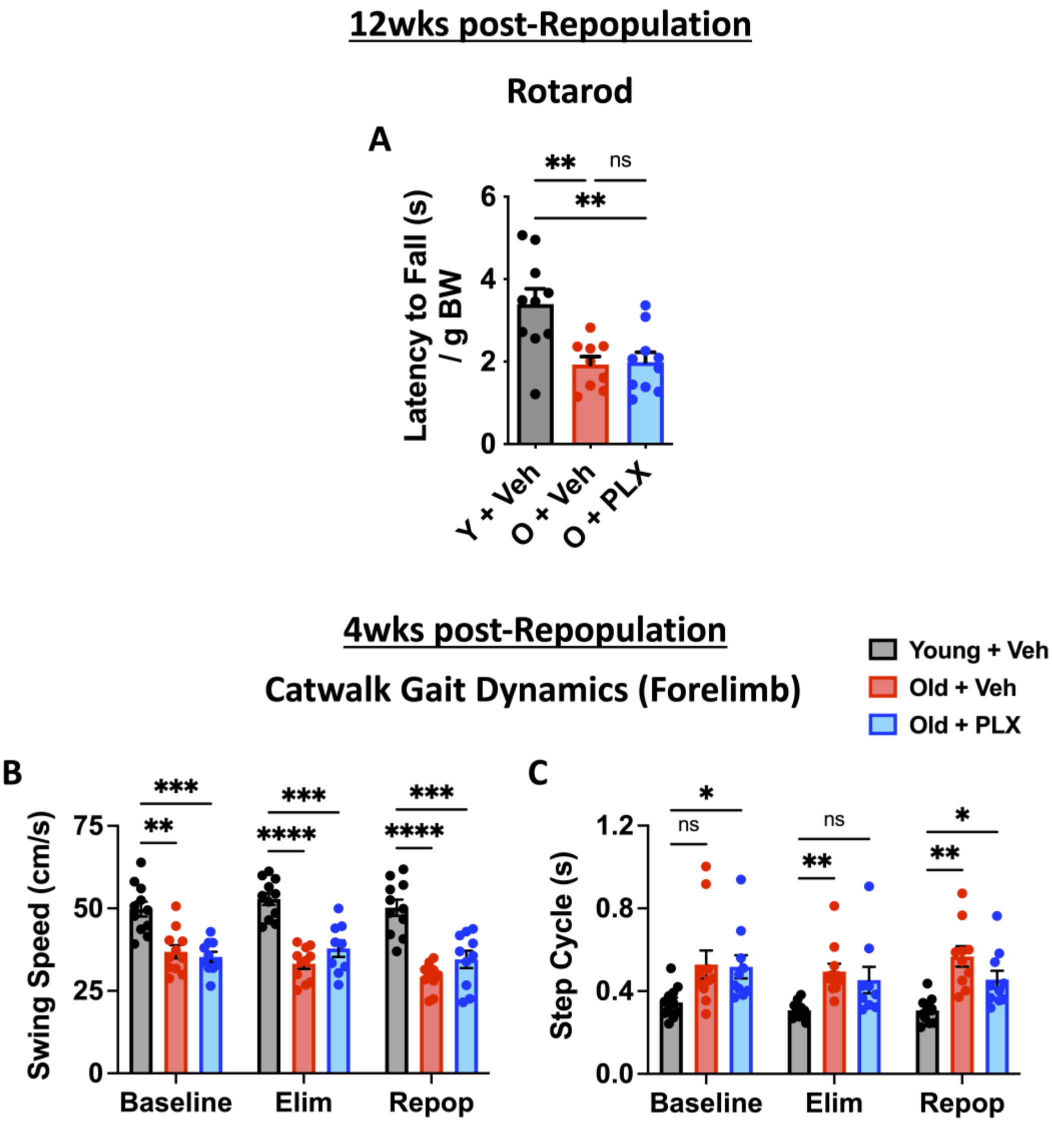


**Fig. S5. Assessment of motor function at 4- and 12-weeks after microglial repopulation.** At 12 weeks post-repopulation, no change in **(A)** rotarod performance was seen. Additional gait measures for forelimb **(B)** swing speed and **(C)** step cycle at four weeks post-repopulation are shown. Abbreviations: cm centimeters, BW body weight, Elim elimination phase, g grams, ns not significant, O old, PLX PLX5622, Repop repopulation phase, s seconds, Veh vehicle, wks weeks, Y young. Data **(B-C)** were analyzed using two-way ANOVA with repeated measures. Data **(A)** were analyzed using one-way ANOVA with multiple comparisons (*p<0.05, **p<0.01, ***p<0.001, and ****p<0.0001).
