## Supplemental Figure 6 for "Brain injury accelerates the onset of a reversible age-related microglial phenotype associated with hyperphagocytosis and inflammatory neurodegeneration"

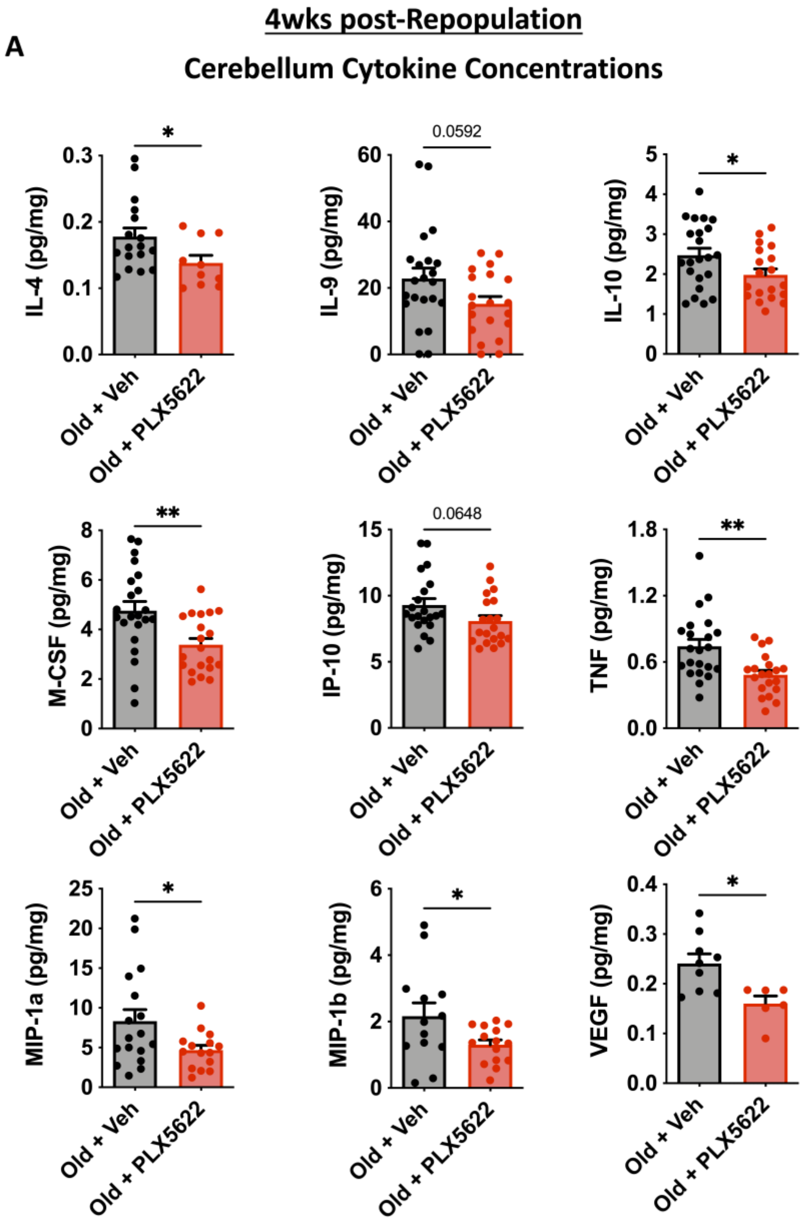


**Fig. S6. Cytokine analysis in the aged cerebellum at four weeks after microglial repopulation.**

Inflammation in the cerebellum of old mice was examined to better understand the treatment effects on age-related motor function. Out of the 32 analytes tested in a multiplex enzyme-linked immunosorbent assay of inflammatory cytokines, those which were detectable and statistically significant or trending after treatment are shown **(A)**. Four weeks after microglial repopulation, cerebellar concentrations of IL-4, IL-10, M-CSF, TNF, MIP-1a, MIP-1b, and VEGF were all significantly reduced compared to the vehicle group. Concentrations of IL-9 and IP-10 showed a non-statistical trend for decrease after treatment. Abbreviations: mg milligram, pg picogram, Veh vehicle. Data **(A)** were analyzed using Student's t-test (*$p<0.05$ and **$p<0.01$).
