## Supplemental Figure 7 for "Brain injury accelerates the onset of a reversible age-related microglial phenotype associated with hyperphagocytosis and inflammatory neurodegeneration"

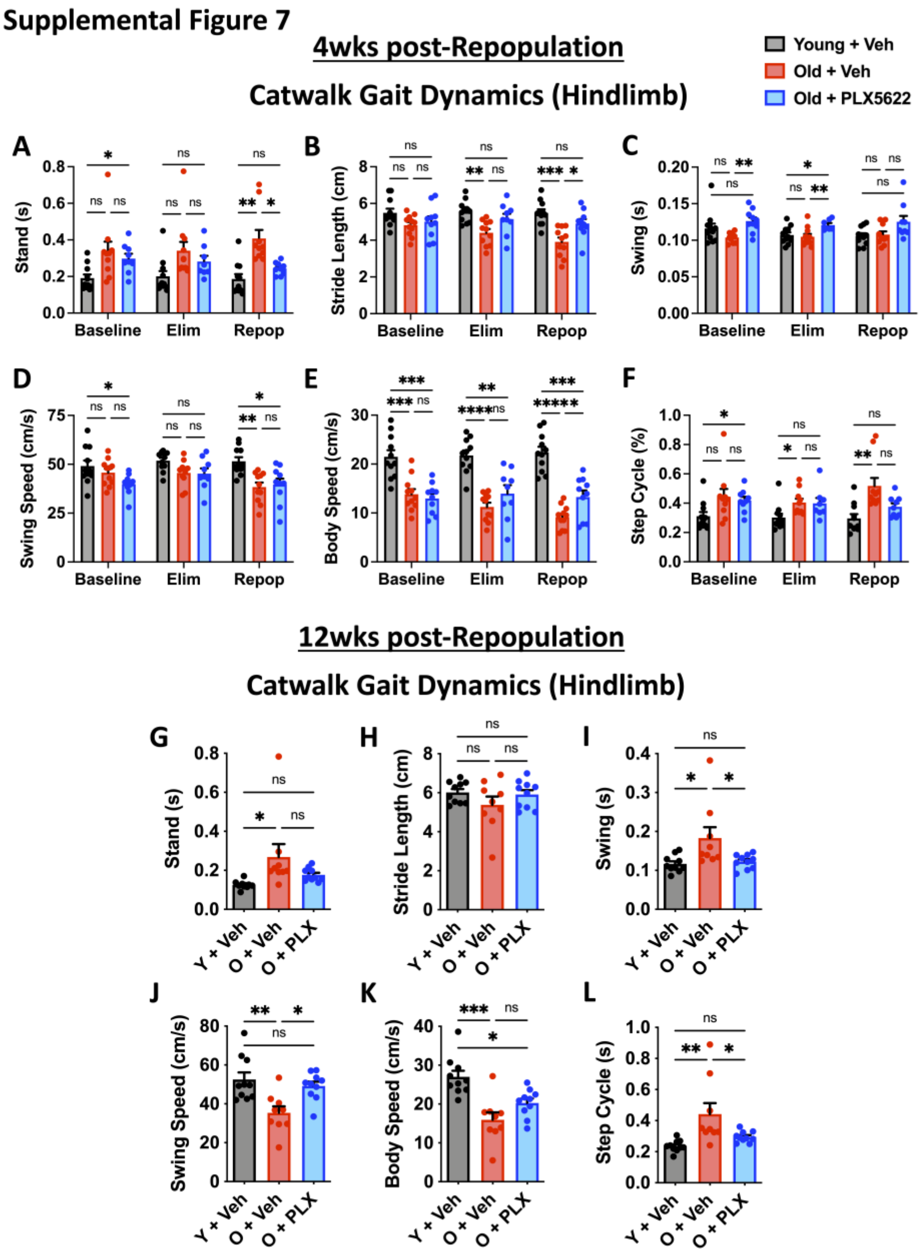


**Fig. S7. Assessment of hindlimb gait dynamics at 4- and 12-weeks after microglial repopulation.**

Gait measures for hindlimb **(A)** stand, **(B)** stride length, **(C)** swing, **(D)** swing speed, **(E)** body speed, and **(F)** step cycle at four weeks post-repopulation are shown. At 12 weeks post-repopulation, hindlimb measures for **(G)** stand, **(H)** stride length, **(I)** swing, **(J)** swing speed, **(K)** body speed, and **(L)** step cycle parameters generally mirror that seen in the forelimbs. Abbreviations: cm centimeters, BW body weight, Elim elimination phase, g grams, ns not significant, O old, PLX PLX5622, Repop repopulation phase, s seconds, Veh vehicle, wks weeks, Y young. Data **(A-F)** were analyzed using two-way ANOVA with repeated measures. Data **(G-L)** were analyzed using one-way ANOVA with multiple comparisons (*$p<0.05$, **$p<0.01$, ***$p<0.001$, and ****$p<0.0001$).
