## Supplemental Figure 8 for "Brain injury accelerates the onset of a reversible age-related microglial phenotype associated with hyperphagocytosis and inflammatory neurodegeneration"

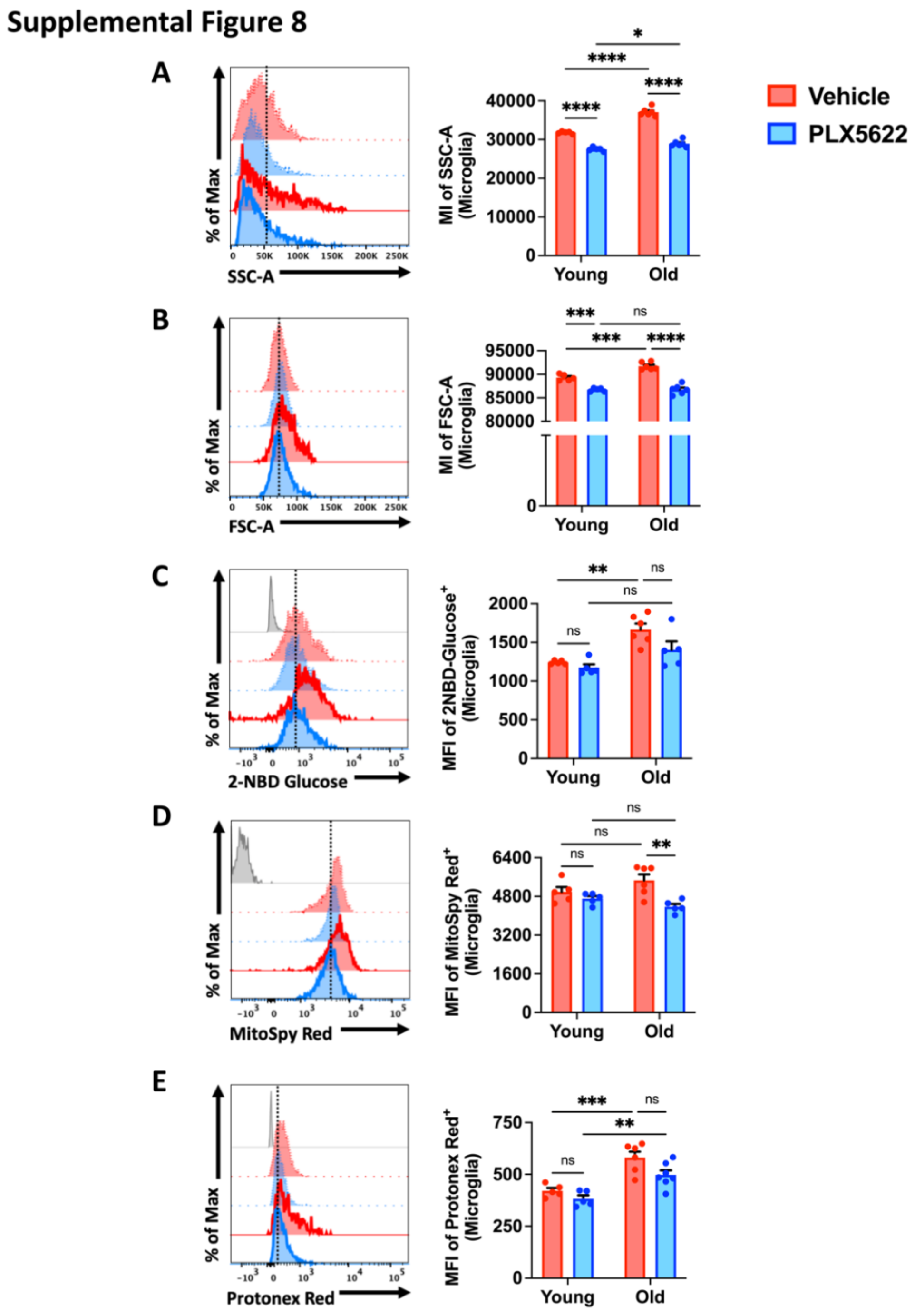


**Fig. S8. Assessment of microglia phenotype at 12 weeks post-repopulation.** Repopulated microglia from old mice treated with PLX5622 displayed significant reductions in **(A)** cell granularity, **(B)** cell volume, **(C)** glucose uptake, **(D)** mitochondrial membrane potential, and **(E)** cytosolic acidosis as evidenced by MFI quantification of the representative histograms. Abbreviations: FSC forward scatter, Max maximum, MFI mean fluorescence intensity, ns not significant, SSC side scatter. Data **(A-E)** were analyzed using two-way ANOVA group analysis with multiple comparison's test (*$p<0.05$, **$p<0.01$, ***$p<0.001$, and ****$p<0.0001$).
