## Supplemental Figure 9 for "Brain injury accelerates the onset of a reversible age-related microglial phenotype associated with hyperphagocytosis and inflammatory neurodegeneration"

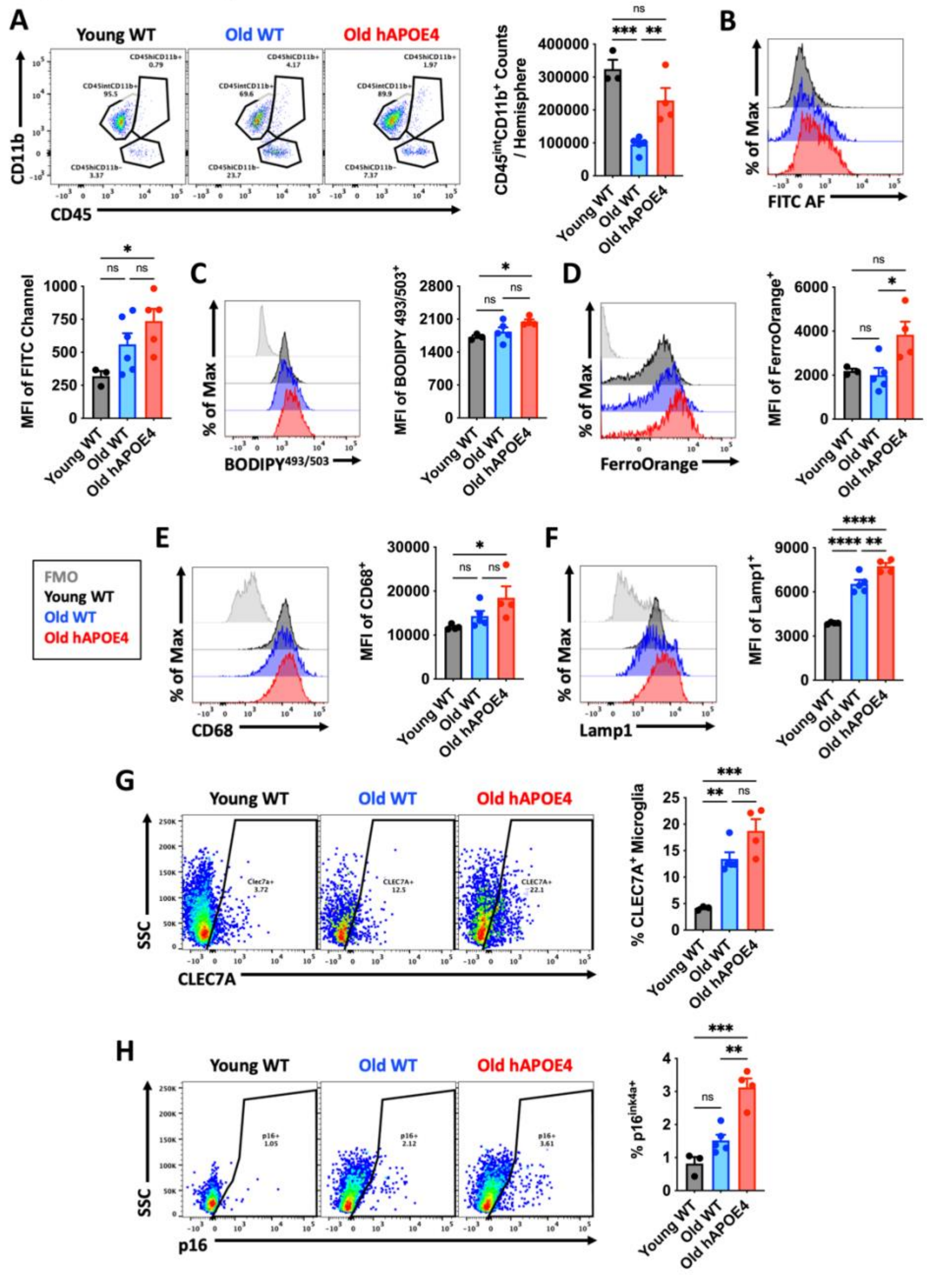


**Fig. S9. Age-associated AF phenotype is modified by hAPOE4 genotype.**
**(A)** A representative dot plot illustrates the immune composition in the brain of young and aged wildtype, and old, age-matched hAPOE$^{+/+}$ knock-in mice. Microglia from aged hAPOE4 mice displayed significantly higher **(B)** autofluorescence, **(C)** lipid content, **(D)** iron concentration, and expression of **(E)** CD68, **(F)** Lamp1, **(G)** CLEC7a, and **(H)** p16$^{ink4a}$ compared to either young or old wildtype controls. Abbreviations: AF autofluorescence, FMO fluorescence minus one control, max Maximum, MFI mean fluorescence intensity, ns not significant, SSC side scatter, WT wildtype. Data **(A-H)** were analyzed using one-way ANOVA with multiple comparisons (N= 3-6/grp; *p<0.05, **p<0.01, ***p<0.001, and ****p<0.0001).
