## Supplemental Figure 10 for "Brain injury accelerates the onset of a reversible age-related microglial phenotype associated with hyperphagocytosis and inflammatory neurodegeneration"

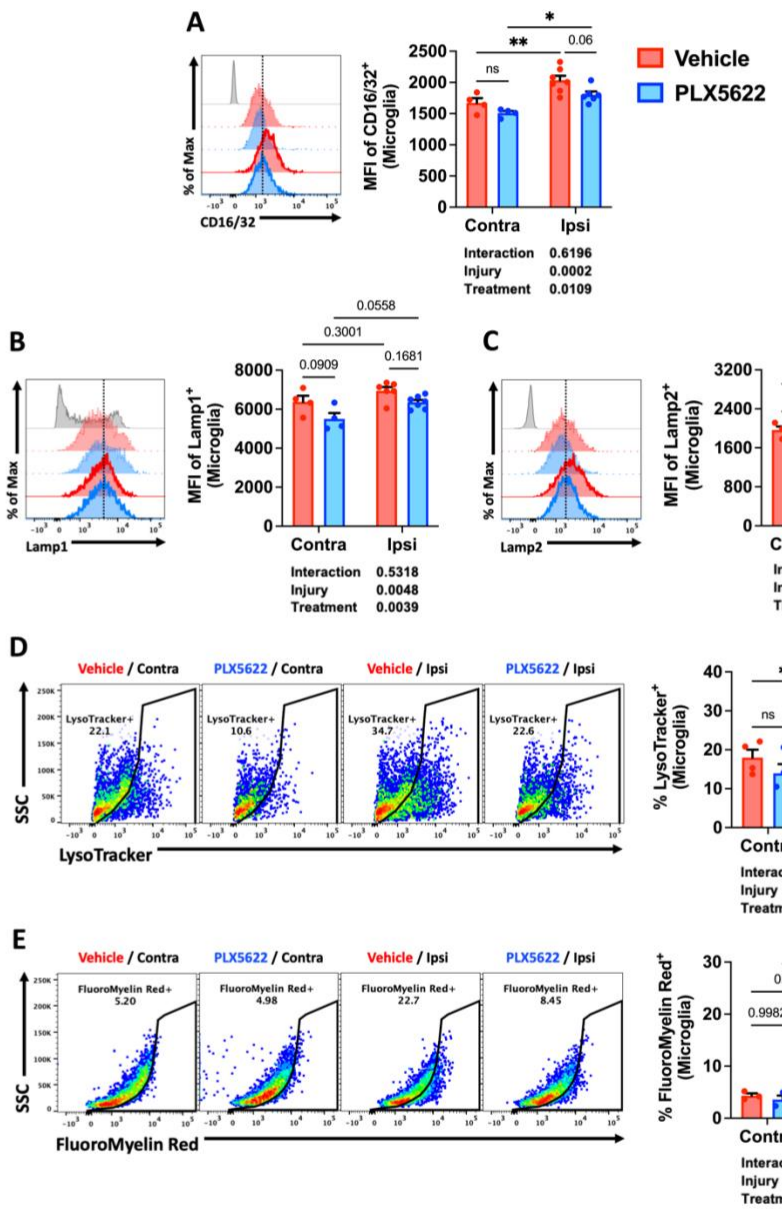


**Fig. S10. Effect of repopulation on microglial phenotype at two weeks post-TBI.** Old mice were treated with vehicle or PLX5622 for two weeks, followed by 12 weeks of repopulation prior to TBI. Microglia in the injured ipsilateral and contralateral control hemisphere were evaluated for phagocytosis and autophagy markers at 2 weeks post-TBI. Representative histograms and relative MFI quantification show that repopulated microglia exhibit treatment effects in **(A)** CD16/32, **(B)** Lamp1, and **(C)** Lamp2 expression compared to the vehicle group after injury. Representative dot plots also demonstrate significant TBI-related reductions in the frequency of **(D)** LysoTracker-positive and **(E)** Fluoromyelin Red-positive microglia. Abbreviations: Contra contralateral, Ipsi ipsilateral, max Maximum, MFI mean fluorescence intensity, ns not significant, SSC side scatter. Data **(A-E)** were analyzed using two-way ANOVA with multiple comparisons (N= 4-7/grp; *$p<0.05$, **$p<0.01$, and ***$p<0.001$).
